# Fast and efficient CRISPR-mediated genome editing in *Aureobasidium pullulans* using Cas9 ribonucleoproteins

**DOI:** 10.1101/2021.07.19.452881

**Authors:** Johanna Kreuter, Georg Stark, Robert L. Mach, Astrid R. Mach-Aigner, Christian Derntl

## Abstract

*Aureobasidium pullulans* is a ubiquitous, polyextremotolerant, “yeast-like” ascomycete used for the industrial production of pullulan and other products and as biocontrol agent in the agriculture. Its application potential and its wide-spread occurrence make *A. pullulans* an interesting study object. The availability of a fast and efficient genome editing method is an obvious advantage for future basic and applied research on *A. pullulans*. In this study, we describe the development of a CRISPR/Cas9-based genome editing method using ribonucleoproteins (RNPs). We demonstrate that this method can be used for single and multiplex genome editing using only RNPs by targeting *ura3* (encoding for orotidine-5′-phosphate decarboxylase), *praics* (encoding for phosphoribosyl aminoimidazole-succinocarboxamide synthase) and *asl* (encoding for arginine succinate lyase). We demonstrate the applicability of *Trichoderma reesei pyr4* and *Aspergillus fumigatus pyrG* to complement the *ura3* deficiency. Further, we show that the usage of RNPs can boost the homologous recombination rate up to nearly 100%, even when using only 20bp long homologous flanks. Therefore, the repair cassettes can be constructed by a single PCR, abolishing the need for laborious and time-consuming cloning. The here presented method allows fast and efficient genome editing for gene deletions, modifications, and insertions in *A. pullulans*.

## Introduction

*Aureobasidium pullulans* is a ubiquitous, black yeast-like ascomycete (*Dothideomycetes*, *Dothideales*), characterized by the production of melanin, phenotypic plasticity, polyextremotolerance and adaptability (Cooke, 1959; de Hoog, 1993; Schoch *et al.*, 2006; Gostinčar *et al.*, 2011). *A. pullulans* is used industrially for the production of pullulan (Bernier, 1958; Bender *et al.*, 1959; Leathers, 2003). Pullulan and its derivatives have a multitude of practical applications in the food, pharmaceutical, agricultural, and chemical industries (Leathers, 2003; Chi *et al.*, 2009). Further products of *A. pullulans* with potential industrial applications are other extracellular polysaccharides, enzymes, antimicrobial compounds, siderophores, heavy oils, poly(β-L-malic acid) (Chi *et al.*, 2009; Prasongsuk *et al.*, 2018). Further, *A. pullulans* can be used as a biocontrol agent in the agriculture sector (Sharma *et al.*, 2009). Based on the wide-spread occurrence and the application potential of *A. pullulans*, there is an obvious demand for an easy and efficient genome editing method.

The clustered regularly interspaced short palindromic repeat (CRISPR) system from *Streptococcus pyogenes* has been used for genome editing in various organisms due to the ease of target programming, modification efficiency, and multiplexing capacity (Doudna and Charpentier, 2014; Sternberg and Doudna, 2015). The modified system depends on a single multifunctional Cas protein (Cas9) and a single guide RNA (sgRNA) which programs Cas9 to introduce a double-strand break (DSB) in a 20 nt-target sequence upstream of a protospacer adjacent motif (PAM, 5’-NGG-3’) (Jinek *et al.*, 2012; Sternberg and Doudna, 2015). Subsequent to the DSB, two main repair pathways i.e., the error-prone non-homologous end joining (NHEJ) and the homology directed repair (HDR) can be exploited for genome editing. The NHEJ repair pathway readily ligates DSBs but often causes insertion/deletion mutations at the target site that can lead to loss of gene function. The HDR pathway can be utilized to insert a defined sequence at the target site. A repair or donor DNA template must be provided to this end (Hsu *et al.*, 2014; Sander and Joung, 2014). There are different methods for delivery of Cas9 and sgRNA into cells available (Yip, 2020). DNA carrying the genes for Cas9 and sgRNA can be transformed. This is cost-effective but requires cloning steps and the plasmid DNA might be inserted at unwanted sites in the genome. Further, the prolonged expression of Cas9 increases the chance of off-target effects (Yip, 2020). Second, the mRNA for the Cas9 can be transformed together with the sgRNA. This minimizes the risk of unwanted integration and off-target effects but is expensive. Third, ribonucleoproteins (RNPs) consisting of the Cas9 protein and the sgRNA can be assembled in vitro and inserted into the target cell. This is a fast and easy delivery technique of CRISPR components that does not require cloning or in vivo transcription and translation. RNPs enable immediate transient gene editing with reduced off-target effects (Kim *et al.*, 2014; Yip, 2020). Cas9-sgRNA RNPs have been shown to efficiently edit the genomes of human and animal cells (Cho *et al.*, 2013; Kim *et al.*, 2014; Chaverra-Rodriguez *et al.*, 2018; Chen *et al.*, 2019), plant cells (Park *et al.*, 2019; Lee *et al.*, 2020) and various fungi (Foster *et al.*, 2018; Zou *et al.*, 2020). In *A. pullulans*, CRISPR mediated genome editing was previously performed using plasmids (Zhang *et al.*, 2019) but RNPS have not yet been used.

In this study, we demonstrate that Cas9-sgRNA RNPs can be used for single and multiplex genome-editing of three *A. pullulans* strains (EXF-150, ATCC 42023 and NBB 7.2.1) by targeting the *ura3* (encoding for orotidine-5′-phosphate decarboxylase), *praics* (encoding for phosphoribosyl aminoimidazole-succinocarboxamide synthase) and *asl* (encoding for arginine succinate lyase) genes. Further, we complemented the uridine auxotrophy with *ura3* homologues from *Trichoderma reesei* and *Aspergillus fumigatus*. Lastly, we demonstrate that integration cassettes with flanks as short as 20 bp can be used for an HDR-mediated gene insertion with homologous integration rates of up to 100%.

## Results and Discussion

### CRISPR/Cas9 RNPs can be used for genome editing in *A. pullulans*

To test, whether Cas9 RNPs can be used in *A. pullulans*, we used the *ura3* gene as a target, because loss-of-function mutations in this gene results in a resistance against 5-fluoroorotic acid (5-FOA) (Rose *et al.*, 2000). We designed two sgRNAs (ura3_sgRNA1 and ura3_sgRNA2) targeting two sites in *ura3* (Fig. 1A). This strategy aimed to enhance the rate of loss-of-function, because the middle gene fragment is expected to get lost during the NHEJ repair. The initial RNP delivery experiments were conducted without sgRNA refolding or the addition of β-mercaptoethanol. For the *A. pullulans* reference strain EXF-150, we obtained about 250 5-FOA resistant colonies after delivering approx. 0.084 nmol of Cas9 and sgRNA each (Fig. S1). To verify that the obtained *ura3* loss-of-function was indeed a result of the CRIPSR mediated DSBs, we sequenced the *ura3* locus of six random colonies. In four colonies (#1, 2, 4, and 5), we observed short deletions at the target site of ura3_sgRNA1 (Fig. 1B), which is a typical result of NHEJ repair mistakes after a DSB (Hsu *et al.*, 2014; Sander and Joung, 2014). In two colonies (#3 and 6), the 800bp-long fragment between the two target sites was deleted (Fig. 1C). Based on this outcome, we conclude that ura3_sgRNA1 is more effective than ura3_sgRNA2. This is in accordance with previous studies; choice of sgRNA affects efficiency and specificity of CRISPR/Cas9 genome editing (Chari, Mali, Moosburner, & Church, 2015; Doench et al., 2016; Doench et al., 2014; Wang, Wei, Sabatini, & Lander, 2014; Xu et al., 2015). However, neither of the six colonies could grow on medium lacking uridine (SC-URA) (Fig. 1D). The uridine auxotrophy could be complemented with the *ura3* homologues from *Trichoderma reesei* (*pyr4*) and *Aspergillus fumigatus* (*pyrG*) (Fig. S2). To this end, the auxotrophic mutant Δ*ura3* #6 was transformed with plasmids pJET-pyr4 (Derntl *et al.*, 2016) and pJET-pyrG, respectively.

**Fig. 1.**
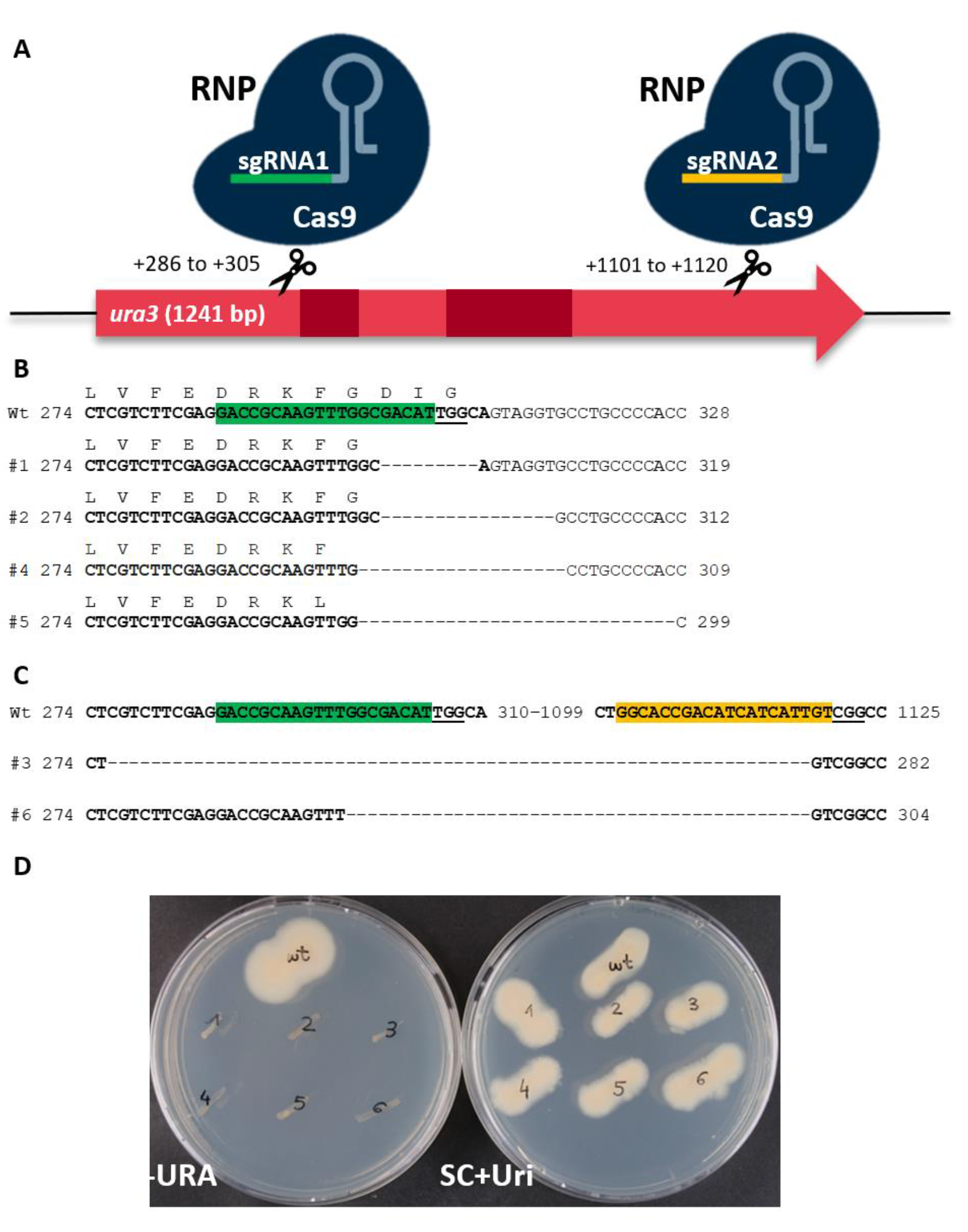
CRISPR/Cas9 mediated manipulation of *ura3* in *A. pullulans* EXF-150 with RNP delivery. **A** Two different sgRNAs (green, ura3_sgRNA1; yellow, ura3_sgRNA2) direct the Cas9 to two distinct target sites in the *ura3* coding region. Introns are indicated in dark red. **B** Partial sequences of *ura3* coding region of the *A. pullulans* EXF-150 wildtype (wt) and six 5-FOA resistant colonies (#1-6) obtained after the delivery of Cas9 RNPs. Exon sequences are bold, the corresponding amino acid sequence is given above the genomic sequence. Target site of ura3_sgRNA1 and ura3_sgRNA2 are highlighted in green and yellow, respectively. PAM sites are underlined. In four colonies (#1, 2, 4 and 5) deletions occurred at the sgRNA1 target site. In #2, 4 and 5, this resulted in a frame shift. In two colonies (#3 and 6) the entire gene fragment between the two target sites (#3: 843 bp segment, #6: 821 bp segment) was lost. **C** The *A. pullulans* EXF-150 wildtype (wt) and six 5-FOA resistant colonies (#1-6) were cultivated on medium lacking uridine (left, SC-URA) and medium containing 5 mM uridine (right, SC+Uri) for 7 days at 24°C.

Next, we tested the applicability of the RNPs in other *A. pullulans* strains. To this end, approx. 0.042 nmol of Cas9 and ura3_sgRNA1 each were delivered into the strains ATCC 42023 and NBB 7.2.1, yielding about 300 and two colonies, respectively (Fig. S3 and Fig. S4**Fehler! Verweisquelle konnte nicht gefunden werden.**). Only ura3_sgRNA1 was used since it was more effective than ura3_sgRNA2 in EXF-150. Sequencing of the *ura3* locus of six random 5-FOA resistant ATCC 42023 colonies and the two 5-FOA resistant NBB 7.2.1 colonies confirmed deletion of nucleotides at the ura3_sgRNA1 target site (Fig. S5 and Fig. S6). Notably, ATCC 42023 was suggested to be *A. pullulans* var. *melanogenum* or *A. melanogenum* in recent studies (Zalar *et al.*, 2008; Rich *et al.*, 2016). Accordingly, we observed an enhanced melanin production in this strain compared to the strains EXF-150 and NBB 7.2.1, and a high sequence similarity of the *ura3* gene to *A. melanogenum* strain TN3-1 (Fig. S7). As the genome of ATCC 42023 is not sequenced, we used ura3_sgRNA1 at a venture and could obtain a relatively high number of colonies (Fig. S3) despite a mismatch in the sgRNA target site (Fig. S5). The low genome editing efficiency of *A. pullulans* NBB 7.2.1 (2 colonies, Fig. S4) could be improved by the addition of β-mercaptoethanol to the protoplasts (as described in (Cullen *et al.*, 1991) and by denaturing and refolding the sgRNA prior to the RNP assembly (as suggested in (Pohl *et al.*, 2018)). These modifications resulted in about 50 colonies for approx. 0.084 nmol Cas9 and sgRNA each (Fig. S8). Consequently, all following RNP delivery approaches were performed with sgRNA denaturing and refolding and β-mercaptoethanol addition.

### Multiplex genome editing allows manipulation of not directly selectable genes

To test for the possibility of multiplex genome editing using Cas9 RNPs, we simultaneously targeted *ura3* and another gene; either the *praics* or the *asl* gene. Loss-of-function mutations in *praics* and *asl* cause adenine and arginine auxotrophy, respectively. For the co-delivery, the sgRNAs targeting *praics* or *asl* were mixed with the ura3_sgRNA1 in a ratio of 11:1 and delivered into *A. pullulans* EXF-150. We selected for *ura3* deficiency (5-FOA resistance) and then tested 24 randomly picked colonies for adenine and arginine auxotrophy, respectively. For *praics*, four out of 24 candidates were adenine auxotroph; they could not grow without adenine (Fig 2A), and turned red, due to the accumulation of the intermediate AIR (5′-phosphoribosyl-5-aminoimidazole) (Fig. 2B). These four candidates carry deletions at the sgRNA target site (Fig. S9). For *asl*, only one out of the tested 24 candidates was arginine auxotroph (Fig. 3), due to mutations at the sgRNA target site (Fig. S10). We speculate that the obtained low frequency of loss-of-function mutations in *praics* and *asl* might be a result of different effectivities of the used sgRNAs. The ura3_sgRNA1 appears to be highly effective. However, the obtained adenine and arginine auxotrophic strains might be used in future studies.

**Fig. 2.**
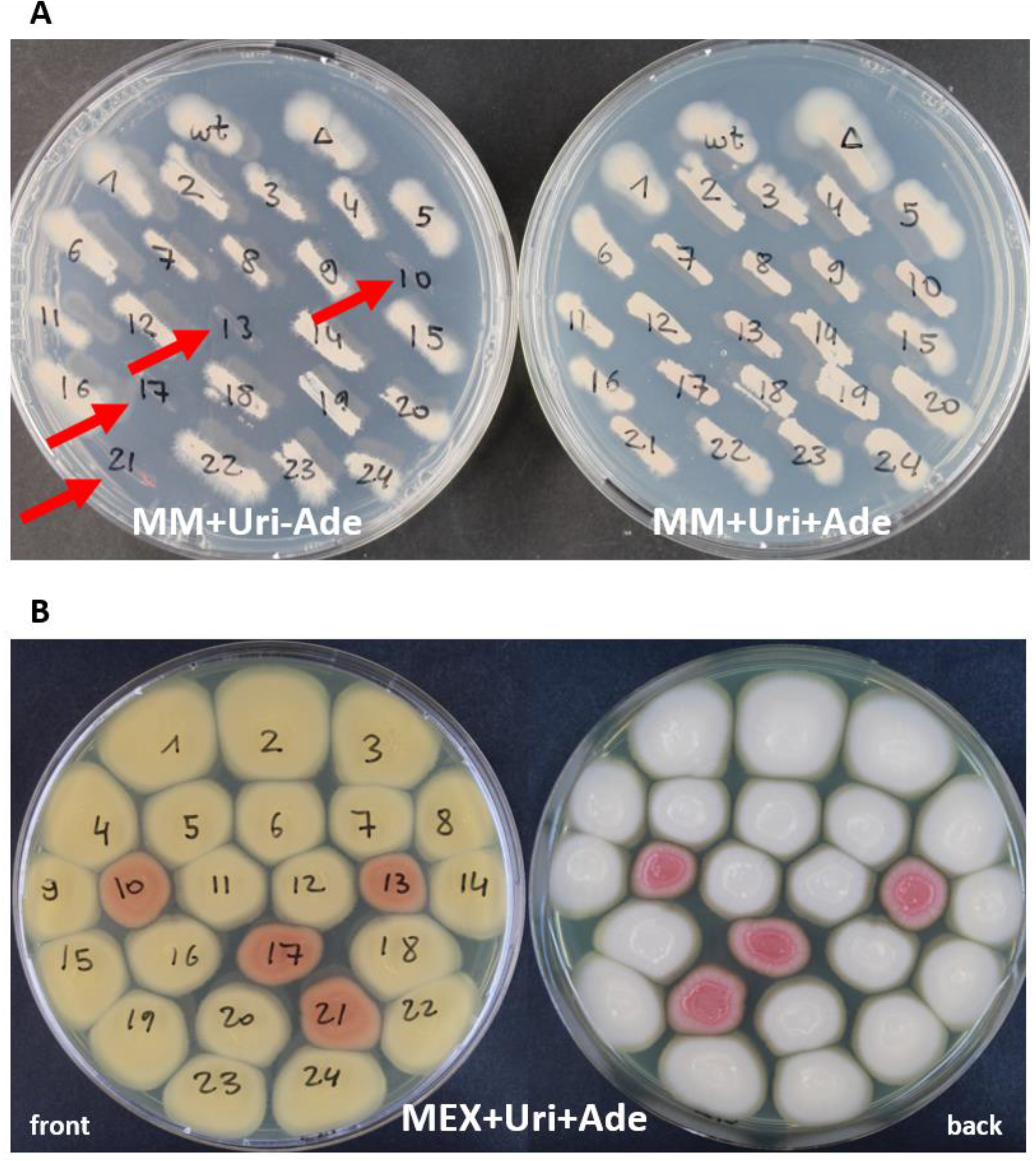
Auxotrophy testing of colonies resulting from a co-delivery of RNPs targeting the *ura3* and the *praic* genes. **A** The *A. pullulans* EXF-150 wildtype (wt), the uridine auxotrophic strain Δura3 #6 (Δ), and 24 randomly selected 5-FOA resistant colonies resulting from the co-delivery of RNPs targeting the *ura3* and the *praics* genes were cultivated on minimal medium lacking adenine (left, MM+Uri-Ade) and on medium containing 0.05 mM adenine (right, MM+Uri+Ade) for 7 days at 24°C. Colonies #10, 13, 17 and 21 were not able to grow on medium without adenine (red arrows). **B** The same strains were cultivated malt extract (MEX) plates supplemented with uridine and adenine (MEX+Uri+Ade) for 7 days at 24°C. Adenine auxotrophic mutants produce a red pigment due to the accumulation of an intermediate in purine biosynthesis.

**Fig. 3.**
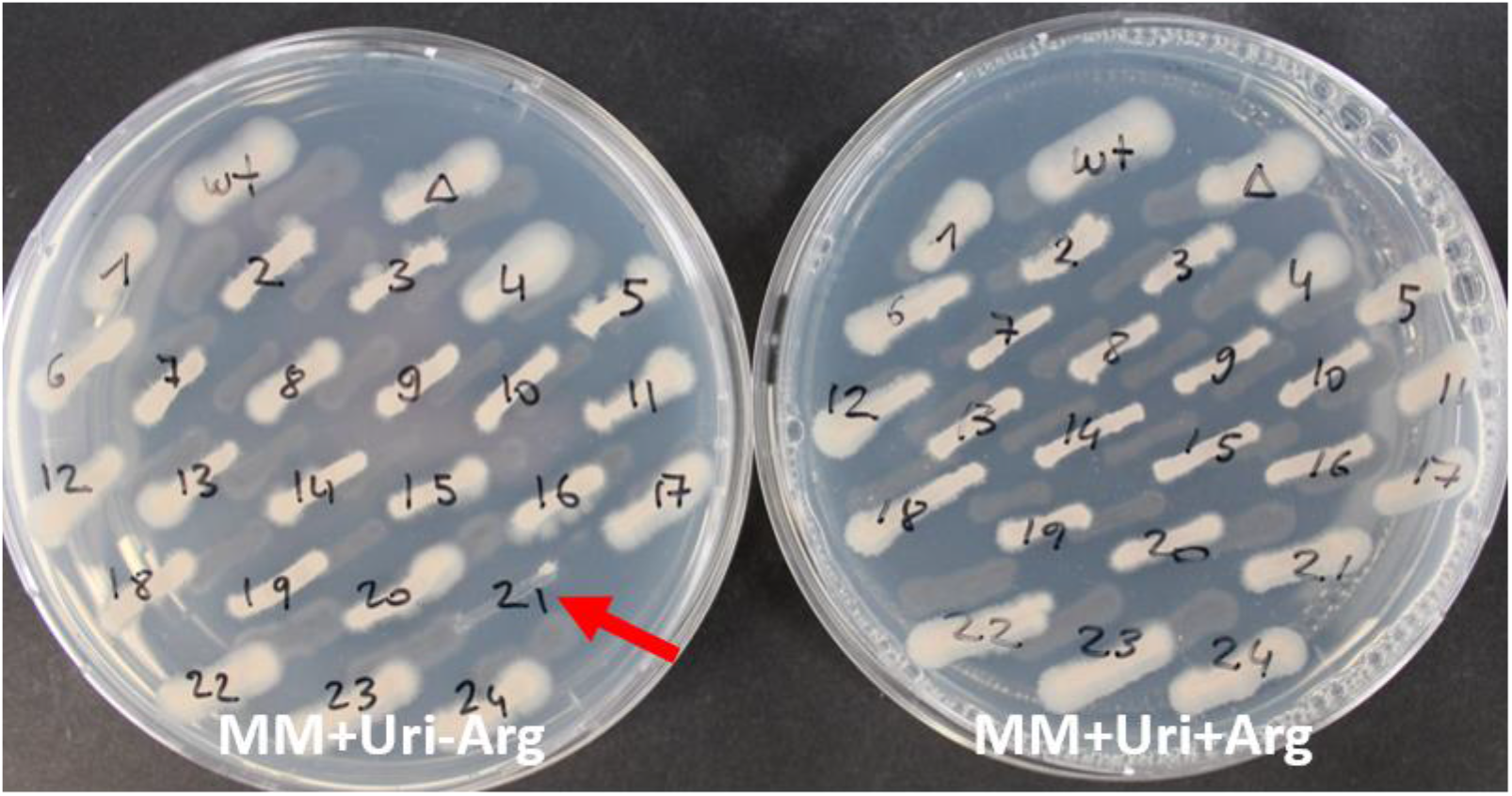
Auxotrophy testing of colonies resulting from a co-delivery of RNPs targeting the *ura3* and the *asl* genes. The *A. pullulans* EXF-150 wildtype (wt), the uridine auxotrophic strain Δura3 #6 (Δ), and 24 randomly selected 5-FOA resistant colonies resulting from the co-delivery of RNPs targeting the *ura3* and the *asl* genes were cultivated on minimal medium lacking arginine (left, MM+Uri-Arg) and medium containing 2.5 mM arginine (right, MM+Uri+Arg) for 7 days at 24°C. Colony #21 was not able to grow on medium without arginine (red arrow).

### Cas9 RNPs can be used to increase the recombination frequency during HDR

Homologous recombination allows advanced genomic manipulations such as the insertion of point mutations, protein tags, and longer genetic material, or exchange of sequences. In many fungi, the NHEJ-pathway is the dominant repair mechanisms and recombination rate is very low (Krappmann, 2007). We were interested whether Cas9 induced DSBs enhance the recombination frequency. To this end, we constructed disruption cassettes and designed a sgRNA targeting the *dl4* gene (encoding DNA ligase IV involved in the NHEJ pathway). The disruption cassettes consisted of the *pyr4* gene from *T. reesei* as marker and homologous flanks of different lengths (20 bp or 500 bp) (Fig. 4). We transformed 3 μg of the disruption cassettes alone or together with dl4_sgRNA-Cas9 RNPs into *A. pullulans* EXF-150 Δ*ura3* #6 and selected for uridine prototrophy. 24 randomly selected colonies for each transformation reaction were tested for homologous integration at the target locus. Without the addition of RNPs, HR frequencies with the 20 bp and 500 bp long flanks were 50% and 83%, respectively (Fig. S11 and Fig. S12). Addition of RNPs increased the frequencies to 96% and 100%, respectively (Fig. S13 and Fig. S14). We verified the homologous recombination after transformation with the 20 bp-long flanks by sequencing (Fig. S15). Homologous recombination could be confirmed in 11 of 12 tested transformants. One transformant showed homologous recombination at the 3’ site but an inconclusive sequencing result at the 5’ site. Notably the corresponding PCR product is longer than expected (Fig. S13). We originally chose this gene with the aim to construct a NHEJ-deficient strain. This strategy has previously been used in a series of fungi to enhance the homologous recombination rate (Krappmann, 2007). To our pleasant surprise, this does not seem to be necessary in *A. pullulans* EXF-150, as the frequency of homologous recombination was already high (50% and 83% using 20 bp and 500 bp-long flanks, respectively). Remarkably, the recombination rate could be enhanced by the usage of RNPs (96% and 100%, respectively). Since 20 bp-long flanks in combination with Cas9 RNPs suffice for a high recombination rate in *A. pullulans* EXF-150, construction of disruption/integration/deletion cassettes can easily be performed via PCR and primers with 20bp-long overhangs.

**Fig. 4.**
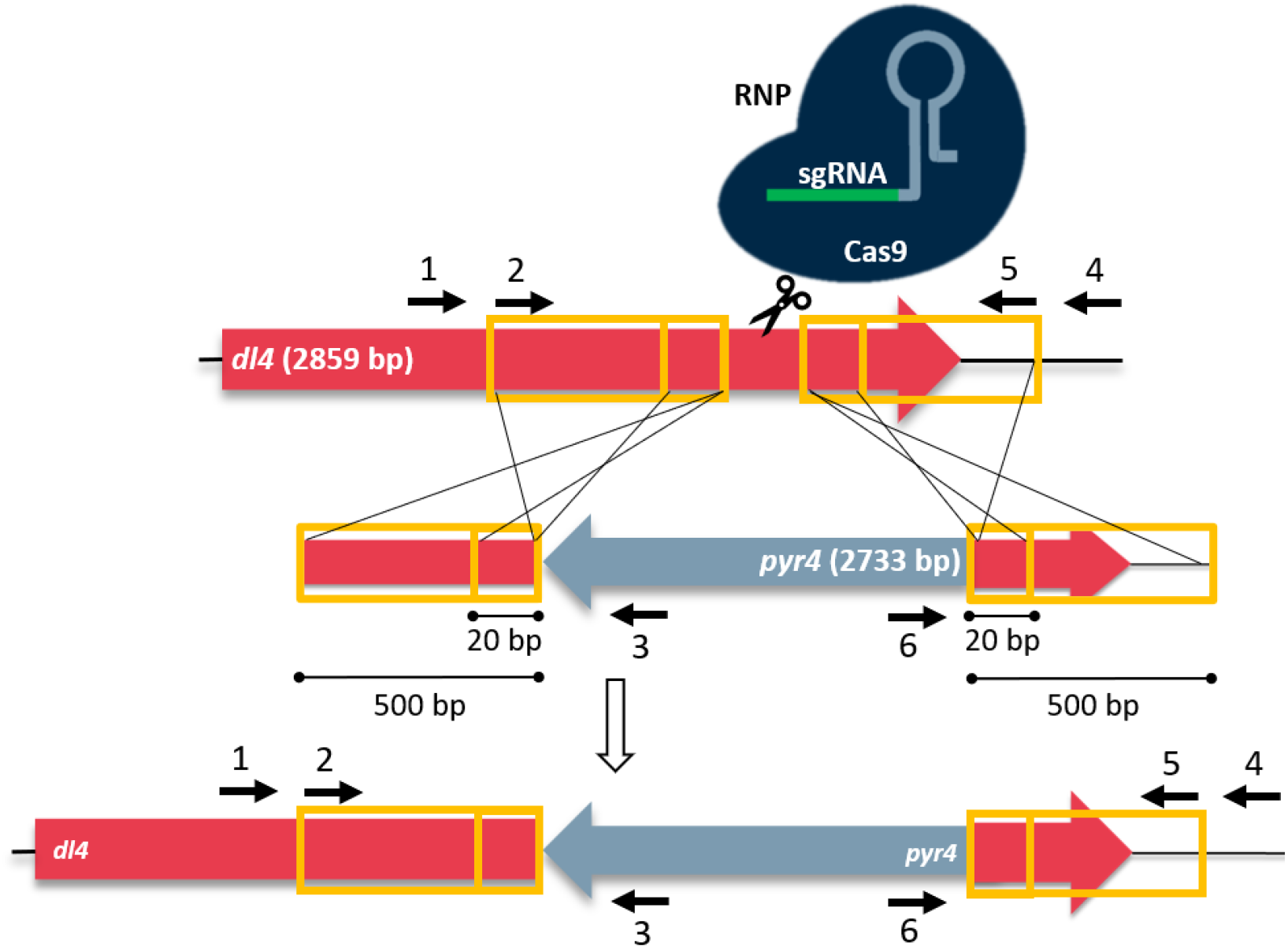
Schematic representation of disruption of *dl4* with *pyr4* disruption cassettes via homologous recombination. The uridine auxotrophic strain (EXF-150 Δura3 #6) was transformed with *pyr4* integration cassettes, in order to complement uridine auxotrophy and disrupt *dl4* (red arrow) with the marker gene *pyr4* (grey arrow) via homologous recombination. The yellow frames represent 5′- and 3′-flanks for the homologous recombination. 20 and 500 bp-long segments in vicinity to the dl4_sgRNA target site were chosen as flanks. The white arrow indicates the recombination event. Primers (black arrows) used for PCR are depicted: 1, 500_flank_fwd; 2, 20_flank_fwd; 3, pyr4_flank_rev; 4, 500_flank_rev; 5, 20_flank_rev; 6, pyr4_flank_fwd. Transformation of donor DNA was carried out with and without delivery of dl4_sgRNA-Cas9 RNPs to assess whether CRISPR/Cas9 RNPs increase HR frequency in *A. pullulans*.

## Experimental Procedures

### Strains and cultivation conditions

*A. pullulans* strains EXF-150 (CBS 100280, (Gostinčar *et al.*, 2014)), ATCC 42023 (Zajic and LeDuy, 1973) and NBB 7.2.1 (CCOS1008, (Hilber-Bodmer *et al.*, 2017)) were maintained on malt extract (MEX) agar at 24°C. Defined medium without yeast extract (Ueda *et al.*, 1963) was used as a minimal medium for testing of adenine and arginine auxotrophy. SC-URA medium (1,71 g l^−1^ Yeast Nitrogen Base, 1,92 g l^−1^ Yeast Synthetic Drop-Out Medium Supplements without Uracil, 5 g l^−1^ (NH_4_)_2_SO_4_ and 20 g l^−1^ glucose) was used as a uridine free medium. If applicable, uridine, 5-fluoroorotic acid (5-FOA), adenine and arginine were added to final concentrations of 5 mM, 2 g l^−1^, 0.5 mM and 2.5 mM, respectively.

### Cas9 protein and sgRNAs

For generation of sgRNAs, target-specific DNA oligonucleotides were designed in silico using the EnGen sgRNA Template Oligo Designer (New England Biolabs, Inc., Ipswich, MA, USA). Templates for sgRNA *in vitro* transcription were synthesized by hybridizing the target-specific oligo and the *S. pyogenes* Cas9 scaffold oligo and filling up with T4 DNA polymerase (New England Biolabs) according to the manufacturer’s instructions. Using this DNA fragment as template, sgRNA was transcribed *in vitro* using the HiScribe Quick T7 High Yield RNA Synthesis Kit (New England Biolabs). The transcribed sgRNA was treated with DNaseI (Thermo Fisher Scientific) and purified using the RNA Cleanup Kit (New England Biolabs). Prior to RNP assembly, the sgRNA was denatured and refolded as described by Pohl et al. (Pohl *et al.*, 2018). RNPs were assembled in a 150 μl reaction in buffer B (1 M sorbitol, 25 mM CaCl_2_, 10 mM Tris.Cl pH 7.5) containing 15 μl 10x Cas9 buffer (20 mM HEPES, 150 mM KCl, 8 mM MgSO_4_· 7 H_2_O, 0.1 mM EDTA, 0.5 mM dithiothreitol, pH 7.5), 4.25 μl EnGen Cas9-NLS (20 μM, New England Biolabs), 2.7 μg sgRNA at 37°C for 10 min.

### RNP delivery and transformation

For the generation of protoplasts 10 ml of an overnight liquid culture with an OD_600_ of approx. 1 were centrifuged at 6000 g for 5 min. The cell pellet was washed with 20 ml buffer A (100 mM KH_2_PO_4_, 1.2 M sorbitol, pH = 5.6) and resuspended in lysing solution (15 ml buffer A containing 150 mg lysing enzymes from *T. harzianum* (Sigma-Aldrich, St. Louis, MO, USA, L1412) and 150 mg β-glucanase from T. *longibrachiatum* (Sigma-Aldrich, G4423). This suspension was incubated at 24°C on a rotary shaker at 140 rpm until protoplasts formed (approx. 1 h). Protoplasts were recovered by the addition of 25 ml ice-cold 1.2 M sorbitol and centrifugation at 4°C and 3000 g for 10 min. Protoplasts were washed once with 30 ml 1.2 M ice-cold sorbitol and twice with 10 ml ice-cold buffer B and then resuspended in 1 ml ice-cold buffer B (volume adjusted to the OD_600_ of the overnight culture). For transformation, 100 μl of the protoplast suspension were mixed with 100 μl “20% PEG solution” (20% (w/v) PEG 4000, 0.67 M sorbitol, 20 mM CaCl2, 10 mM Tris pH = 7.5) and 2 μl β-mercaptoethanol added. Next, 150 μl of the RNP mix or 150 μl Buffer B were added. For transformation of DNA, 5 μg of undigested plasmid DNA or 3 μg of linear donor DNA were used, respectively. The reactions were incubated on ice for 30 minutes and 750 μl “60% PEG solution” (60% (w/v) PEG 4000, 10 mM CaCl2, 10 mM Tris pH = 7.5) added stepwise. After 20 minutes at 23°C, 4.1 ml of buffer C (1 M sorbitol, 10 mM Tris.Cl pH= 7.5) were added stepwise. Different amounts of the transformation mix were added to 20 mL of melted, 50°C warm selection medium, containing 1 M sucrose. This mixture was poured into sterile petri dishes. The plates were incubated at 24°C for 6 to 14 days until colonies were visible.

### Construction of pJET-pyrG

The *pyrG* gene of *A. fumigatus* was amplified by PCR using the Q5 DNA Polymerase (New England Biolabs), the primers pyrG_fwd-AflII-NsiI and pyrG_rev-EcoRI-AatII, and the plasmid pFC330 (Nødvig *et al.*, 2015) as template. The PCR was inserted into pJET1.2 using the CloneJET PCR Cloning Kit (Thermo Scientific) according to the manufacturer’s instructions. The sequence was verified by Sanger sequencing (at Microsynth AG, Balgach, Switzerland) (Fig. S16 and Fig. S17).

### Construction of disruption cassettes

For the construction of pUC18_Apdp4 (Fig. S18) a gene assembly strategy using the NEBuilder HiFi DNA Assembly Cloning Kit (New England Biolabs) was followed. The *pyr4* gene was amplified using the primers pyr4dl4_20bp_rev and pyr4dl4_20bp_fwd and the plasmid pJET-pyr4 (Derntl *et al.*, 2016) as template. The two 500 bp-long homology flanks of the *dl4* gene were amplified using the primers dl4_5Overlap500_fwd and dl4_5Overlap_rev or dl4_3Overlap_fwd and dl4_3Overlap500_rev and genomic DNA of *A. pullulans* EXF-150 as template. The plasmid pUC18 was amplified with the primers pUC18_fwd and pUC18_rev. The sequence of pUC18_Apdp4 was verified by Sanger sequencing (at Microsynth AG). Linear disruption cassettes (Fig. S19) for transformation were amplified using the primers dl4_20bpover_fwd and dl4_20bpover_rev or dl4_500bpover_fwd and dl4_500bpover_rev and pUC18_Apdp4 as template. The PCR products were purified with the GeneJet PCR Purification Kit (Thermo Scientific) according to the manufacturer’s instructions. Several PCR reactions were pooled to obtain enough DNA for transformation. The DNA was concentrated by precipitation with sodium acetate and ethanol and dissolved in double-distilled water (ddH_2_O).

### Genotyping

For the extraction of chromosomal DNA, we used the colony PCR protocol described by Wu et al. (Wu *et al.*, 2017). For diagnostic PCR, 2μL of the resulting crude DNA extract was used as the template in a 50-μl PCR with the OneTaq DNA polymerase (New England Biolabs) according to the manufacturer’s instructions. For subsequent agarose gel electrophoresis of the DNA fragments, a GeneRuler 1-kb Plus DNA ladder (New England Biolabs) was applied to estimate the fragment size. Loci to be sequenced were amplified by PCR with the Q5 DNA polymerase (New England Biolabs) according to the manufacturer’s instructions and sequenced at Microsynth AG.

### Oligonucleotides

All oligonucleotides used in this study are listed in Table S1.

## Acknowledgements

This study was supported by the Austrian Science Fund (FWF, https://www.fwf.ac.at/) [P 29556 to RM, P 34036 to CD]. The authors declare that is no conflict of interest.

## Notes

### Competing Interest Statement

The authors have declared no competing interest.

## References

Bender, H., Lehmann, J., and Wallenfels, K. (1959) Pullulan, ein extracelluläres Glucan von *Pullularia pullulans*, Biochimica et Biophysica Acta 36: 309–316.

Bernier, B. (1958) THE PRODUCTION OF POLYSACCHARIDES BY FUNGI ACTIVE IN THE DECOMPOSITION OF WOOD AND FOREST LITTER, Canadian Journal of Microbiology 4: 195–204.

Branda, E., Turchetti, B., Diolaiuti, G., Pecci, M., Smiraglia, C., and Buzzini, P. (2010) Yeast and yeast-like diversity in the southernmost glacier of Europe (Calderone Glacier, Apennines, Italy), FEMS Microbiology Ecology 72: 354–369.

Chaverra-Rodriguez, D., Macias, V.M., Hughes, G.L., Pujhari, S., Suzuki, Y., Peterson, D.R. et al. (2018) Targeted delivery of CRISPR-Cas9 ribonucleoprotein into arthropod ovaries for heritable germline gene editing, Nature Communications 9: 3008.

Chen, S., Sun, S., Moonen, D., Lee, C., Lee, A.Y.-F., Schaffer, D.V., and He, L. (2019) CRISPR-READI: Efficient Generation of Knockin Mice by CRISPR RNP Electroporation and AAV Donor Infection, Cell Rep 27: 3780–3789.e3784.

Chi, Z., Wang, F., Chi, Z., Yue, L., Liu, G., and Zhang, T. (2009) Bioproducts from *Aureobasidium pullulans*, a biotechnologically important yeast, Applied Microbiology and Biotechnology 82: 793–804.

Cho, S.W., Lee, J., Carroll, D., Kim, J.-S., and Lee, J. (2013) Heritable gene knockout in *Caenorhabditis elegans* by direct injection of Cas9-sgRNA ribonucleoproteins, Genetics 195: 1177–1180.

Cooke, W.B. (1959) An ecological life history of *Aureobasidium pullulans* (de Bary) Arnaud, Mycopathologia et mycologia applicata 12: 1–45.

Cullen, D., Yang, V., Jeffries, T., Bolduc, J., and Andrews, J.H. (1991) Genetic transformation of *Aureobasidium pullulans*, Journal of Biotechnology 21: 283–288.

de Garcia, V., Brizzio, S., and van Broock, M.R. (2012) Yeasts from glacial ice of Patagonian Andes, Argentina, FEMS Microbiology Ecology 82: 540–550.

de Hoog, G.S. (1993) Evolution of black yeasts: possible adaptation to the human host, Antonie van Leeuwenhoek 63: 105–109.

Derntl, C., Rassinger, A., Srebotnik, E., Mach, R.L., and Mach-Aigner, A.R. (2016) Identification of the Main Regulator Responsible for Synthesis of the Typical Yellow Pigment Produced by *Trichoderma reesei*, Appl Environ Microbiol 82: 6247–6257.

Doudna, J.A., and Charpentier, E. (2014) The new frontier of genome engineering with CRISPR-Cas9, Science 346: 1258096.

Foster, A.J., Martin-Urdiroz, M., Yan, X., Wright, H.S., Soanes, D.M., and Talbot, N.J. (2018) CRISPR-Cas9 ribonucleoprotein-mediated co-editing and counterselection in the rice blast fungus, Scientific Reports 8: 14355.

Gostinčar, C., Grube, M., and Gunde-Cimerman, N. (2011) Evolution of Fungal Pathogens in Domestic Environments?, Fungal Biology 115: 1008–1018.

Gostinčar, C., Ohm, R.A., Kogej, T., Sonjak, S., Turk, M., Zajc, J. et al. (2014) Genome sequencing of four *Aureobasidium pullulans* varieties: biotechnological potential, stress tolerance, and description of new species, BMC Genomics 15: 549.

Gunde-Cimerman, N., Zalar, P., de Hoog, S., and Plemenitaš, A. (2000) Hypersaline waters in salterns – natural ecological niches for halophilic black yeasts, FEMS Microbiology Ecology 32: 235–240.

Hilber-Bodmer, M., Schmid, M., Ahrens, C.H., and Freimoser, F.M. (2017) Competition assays and physiological experiments of soil and phyllosphere yeasts identify *Candida subhashii* as a novel antagonist of filamentous fungi, BMC Microbiology 17: 4.

Hsu, Patrick D., Lander, Eric S., and Zhang, F. (2014) Development and Applications of CRISPR-Cas9 for Genome Engineering, Cell 157: 1262–1278.

Jinek, M., Chylinski, K., Fonfara, I., Hauer, M., Doudna, J.A., and Charpentier, E. (2012) A programmable dual-RNA-guided DNA endonuclease in adaptive bacterial immunity, Science 337: 816–821.

Kim, S., Kim, D., Cho, S.W., Kim, J., and Kim, J.-S. (2014) Highly efficient RNA-guided genome editing in human cells via delivery of purified Cas9 ribonucleoproteins, Genome Res 24: 1012–1019.

Krappmann, S. (2007) Gene targeting in filamentous fungi: the benefits of impaired repair, Fungal Biology Reviews 21: 25–29.

Leathers, T.D. (2003) Biotechnological production and applications of pullulan, Applied Microbiology and Biotechnology 62: 468–473.

Lee, M.H., Lee, J., Choi, S.A., Kim, Y.-S., Koo, O., Choi, S.H. et al. (2020) Efficient genome editing using CRISPR–Cas9 RNP delivery into cabbage protoplasts via electro-transfection, Plant Biotechnology Reports 14: 695–702.

Li, H., Chi, Z., Wang, X., Duan, X., Ma, L., and Gao, L. (2007) Purification and characterization of extracellular amylase from the marine yeast *Aureobasidium pullulans* N13d and its raw potato starch digestion, Enzyme and Microbial Technology 40: 1006–1012.

Niehaus, F., Bertoldo, C., Kähler, M., and Antranikian, G. (1999) Extremophiles as a source of novel enzymes for industrial application, Applied Microbiology and Biotechnology 51: 711–729.

Nødvig, C.S., Nielsen, J.B., Kogle, M.E., and Mortensen, U.H. (2015) A CRISPR-Cas9 System for Genetic Engineering of Filamentous Fungi, PLOS ONE 10: e0133085.

Park, J., Choi, S., Park, S., Yoon, J., Park, A.Y., and Choe, S. (2019) DNA-Free Genome Editing via Ribonucleoprotein (RNP) Delivery of CRISPR/Cas in Lettuce. In: Plant Genome Editing with CRISPR Systems: Methods and Protocols. Qi, Y. (ed). New York, NY: Springer New York. 337–354.

Pohl, C., Mózsik, L., Driessen, A., Bovenberg, R., and Nygård, Y. (2018) Genome Editing in *Penicillium chrysogenum* Using Cas9 Ribonucleoprotein Particles, Methods in molecular biology 1772: 213–232.

Prasongsuk, S., Lotrakul, P., Ali, I., Bankeeree, W., and Punnapayak, H. (2018) The current status of *Aureobasidium pullulans* in biotechnology, Folia Microbiologica 63: 129–140.

Rich, J.O., Manitchotpisit, P., Peterson, S.W., Liu, S., Leathers, T.D., and Anderson, A.M. (2016) Phylogenetic classification of *Aureobasidium pullulans* strains for production of feruloyl esterase, Biotechnology Letters 38: 863–870.

Rose, K., Liebergesell, M., and Steinbüchel, A. (2000) Molecular analysis of the *Aureobasidium pullulans ura3* gene encoding orotidine-5′-phosphate decarboxylase and isolation of mutants defective in this gene, Applied Microbiology and Biotechnology 53: 296–300.

Sander, J.D., and Joung, J.K. (2014) CRISPR-Cas systems for editing, regulating and targeting genomes, Nat Biotechnol 32: 347–355.

Schoch, C.L., Shoemaker, R.A., Seifert, K.A., Hambleton, S., Spatafora, J.W., and Crous, P.W. (2006) A multigene phylogeny of the *Dothideomycetes* using four nuclear loci, Mycologia 98: 1041–1052.

Sharma, R.R., Singh, D., and Singh, R. (2009) Biological control of postharvest diseases of fruits and vegetables by microbial antagonists: A review, Biol Control 50: 205–221.

Sternberg, Samuel H., and Doudna, Jennifer A. (2015) Expanding the Biologist’s Toolkit with CRISPR-Cas9, Molecular Cell 58: 568–574.

Ueda, S., Fujita, K., Komatsu, K., and Nakashima, Z.I. (1963) Polysaccharide produced by the genus *Pullularia*. I. Production of polysaccharide by growing cells, Appl Microbiol 11: 211–215.

Wu, Y., Li, B.-Z., Zhao, M., Mitchell, L.A., Xie, Z.-X., Lin, Q.-H. et al. (2017) Bug mapping and fitness testing of chemically synthesized chromosome X, Science 355: eaaf4706.

Yip, B.H. (2020) Recent Advances in CRISPR/Cas9 Delivery Strategies, Biomolecules 10: 839.

Zajic, J.E., and LeDuy, A. (1973) Flocculant and chemical properties of a polysaccharide from *Pullularia pullulans*, Appl Microbiol 25: 628–635.

Zalar, P., Gostincar, C., de Hoog, G.S., Ursic, V., Sudhadham, M., and Gunde-Cimerman, N. (2008) Redefinition of *Aureobasidium pullulans* and its varieties, Stud Mycol 61: 21–38.

Zhang, Y., Feng, J., Wang, P., Xia, J., Li, X., and Zou, X. (2019) CRISPR/Cas9-mediated efficient genome editing via protoplast-based transformation in yeast-like fungus *Aureobasidium pullulans*, Gene 709: 8–16.

Zhdanova, N.N., Zakharchenko, V.A., Vember, V.V., and Nakonechnaya, L.T. (2000) Fungi from Chernobyl: mycobiota of the inner regions of the containment structures of the damaged nuclear reactor, Mycological Research 104: 1421–1426.

Zou, G., Xiao, M., Chai, S., Zhu, Z., Wang, Y., and Zhou, Z. (2020) Efficient genome editing in filamentous fungi via an improved CRISPR-Cas9 ribonucleoprotein method facilitated by chemical reagents, Microbial Biotechnology n/a.

