## Supplementary data for "Fast and efficient CRISPR-mediated genome editing in *Aureobasidium pullulans* using Cas9 ribonucleoproteins"

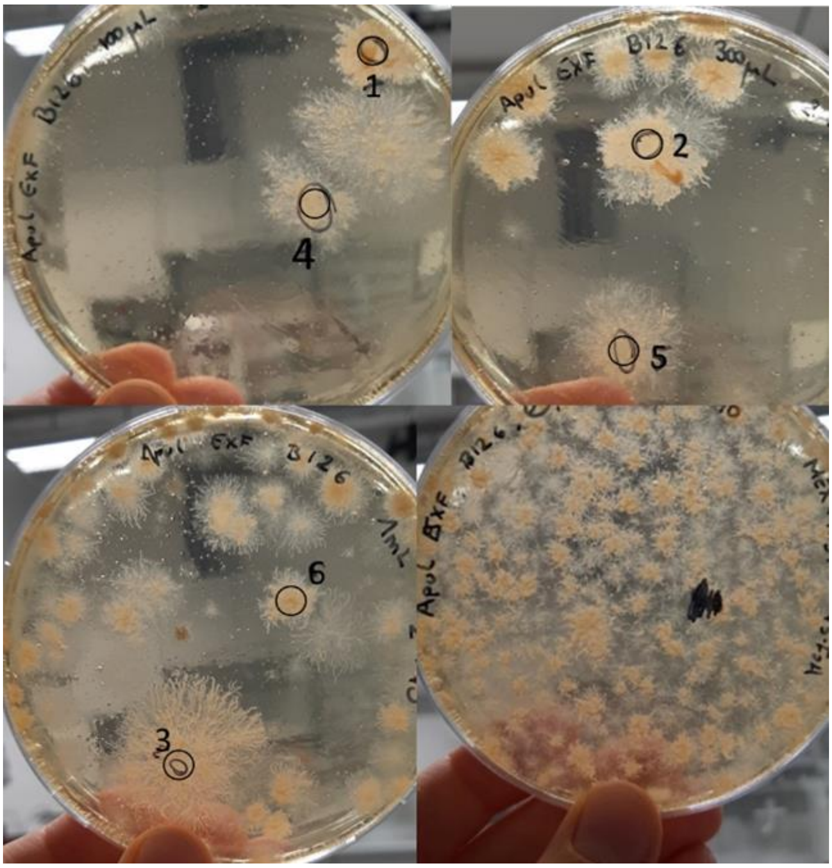


**Fig. S1** Selection plates (MEX + Uri + 5-FOA) resulting from the delivery of ura3_sgRNA1 and ura3_sgRNA2-Cas9 RNPs into *A. pullulans* EXF-150 after 14 days. About 250 CFUs were obtained after the delivery of approx. 0.084 nmol Cas9 and sgRNA each.

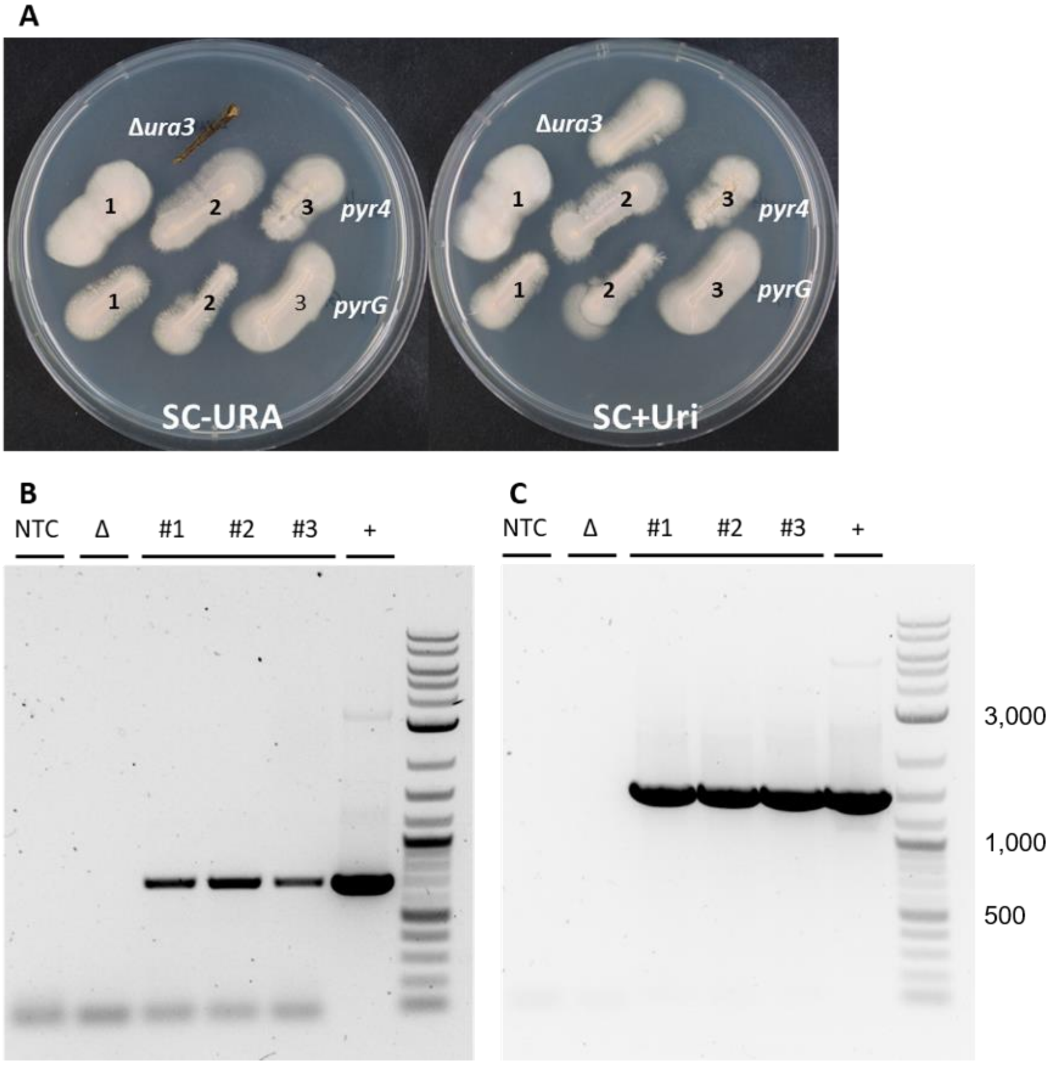


**Fig. S2** Complementation of uridine auxotrophy in *A. pullulans EXF*-150 uridine auxotrophic mutant #6 (Δura3 #6) with *pyr4* (*T. reesei*) and *pyrG* (*A. fumigatus*), respectively.

**A** The parent strain Δura3 #6 cannot grow on medium lacking uridine (SC-URA) (left). Growth is only possible after the addition of 5 mM uridine (SC+Uri) (right). Colonies resulting from the transformation with pJET-pyr4 / pJET-pyrG are able to grow on SC-URA. Pictures taken after 7 days incubation at 24°C.

**B** Complementation of uridine auxotrophy with *pyr4*: Agarose gel electrophoresis of the fragments obtained by PCR with primers pyr4_fwd and pyr4_rev using chromosomal DNA of the parent strain (Δura3 #6, Δ) and candidates (pyr4 #1-3) as template. NTC, no template control; +, positive control pJET-pyr4.

**C** Complementation of uridine auxotrophy with *pyrG*: Agarose gel electrophoresis of the fragments obtained by PCR with primers pyrG_fwd and pyrG_rev using chromosomal DNA of the parent strain (Δura3 #6, Δ) and candidates (pyrG #1-3) as template. NTC, no template control; +, positive control pJET-pyrG.

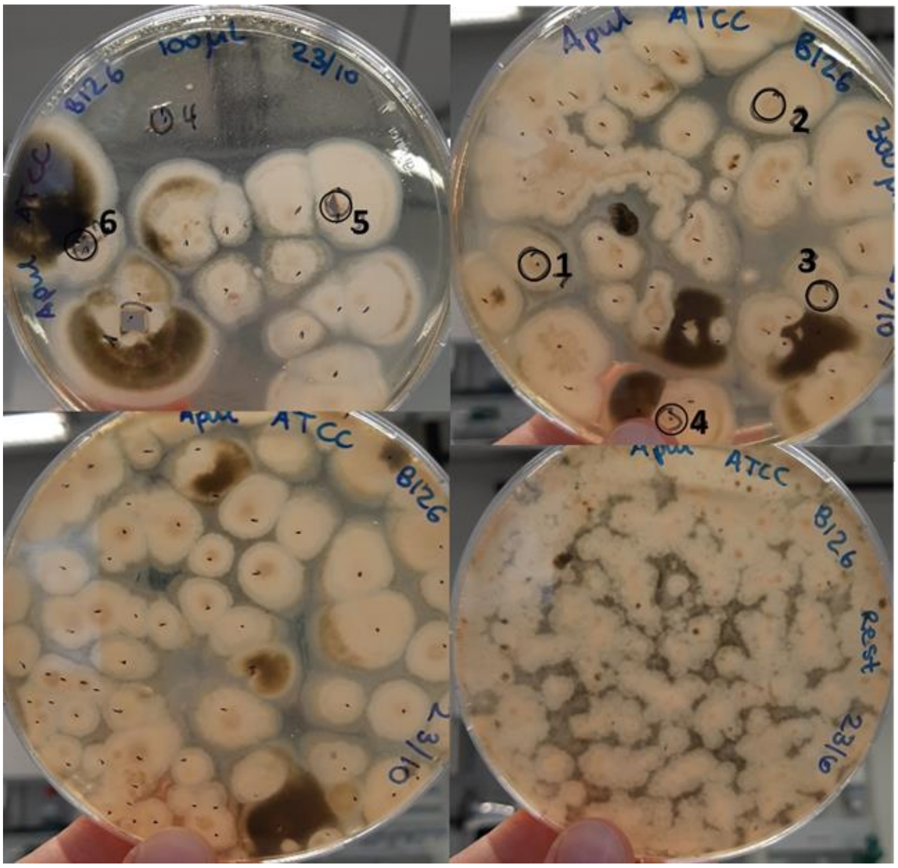


**Fig. S3** Selection plates (MEX + Uri + 5-FOA) resulting from the delivery of ura3_sgRNA1-Cas9 RNPs into *A. pullulans* ATCC 42023 after 11 days. About 300 CFUs were obtained after the delivery of approx. 0.042 nmol Cas9 and sgRNA each.

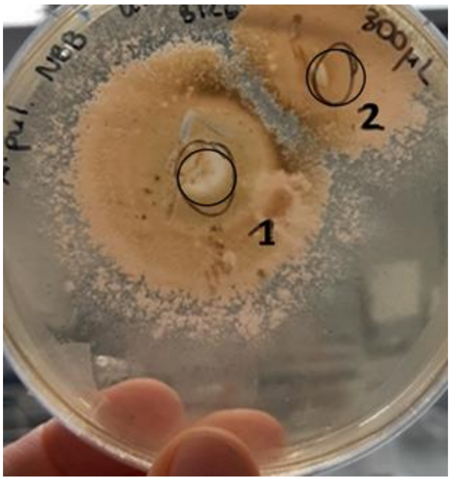


**Fig. S4** Selection plate (MEX + Uri + 5-FOA) resulting from the delivery of ura3_sgRNA1-Cas9 RNPs into *A. pullulans* NBB 7.2.1 after 28 days. Two CFUs were obtained after the delivery of approx. 0.042 nmol Cas9 and sgRNA each.

**A**

```
Wt      GCCAAAAAACACTTCCTCGTTTTCGAGGATCGCAAGTTTGGCGACATTGGCAGTATGTTCATGCCCCACT
#1      GCCAAAAAACACTTCCTCGTTTTCGAGGATCGCAAGTTT--------------TATGTTCATGCCCCACT
#2      GCCAAAAAACACTTCCTCGTTTTCGAGGATCGCAAGTTT--------------TATGTTCATGCCCCACT
#3      GCCAAAAAACACTTCCTCGTTTTCGAGGATCGCAAGTTT---------------ATGTTCATGCCCCACT
#4      GCCAAAAAACACTTCCTCGTTTTCGAGGATCGCCCATT----------GGCAGTATGTTCATGCCCCACT
#5      GCCAAAAAACACTTCCTCGTTTTCGAGGATCGCAAGTTTG------------GAATGTTCATGCCCCACT
#6      GCCAAAAAACACTTCCTCGTTTTCGAGGATCGCAAGTTTGGCG----------TATGTTCATGCCCCACT
```

**B**

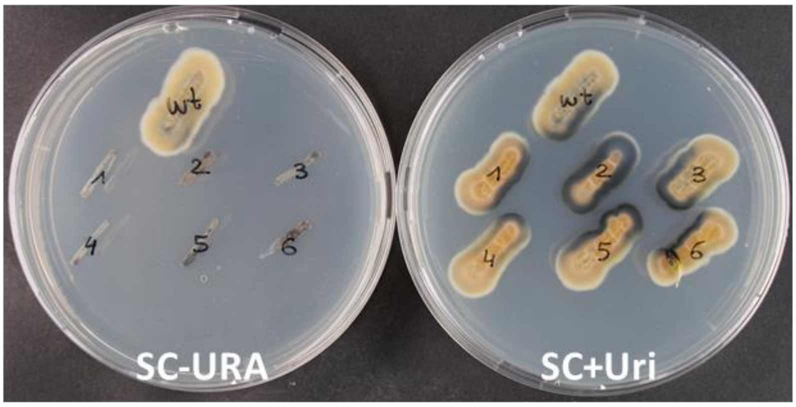


**Fig. S5** CRISPR/Cas9 mediated manipulation of *ura3* in *A. pullulans* ATCC 42023 with RNP delivery.

**A** Partial sequences of *ura3* coding region of the *A. pullulans* ATCC 42023 wildtype (wt) and six 5-FOA resistant colonies (#1-6) obtained after the delivery of Cas9 RNPs. Target site of ura3_sgRNA1 is highlighted in green. Mismatch against the sgRNA is indicated in bold. PAM site is underlined. In all transformants deletions occurred at the sgRNA target site, resulting in a frame shift in colonies #1, 2, 4 and 6.

**B** The *A. pullulans* ATCC 42023 wildtype (wt) and the six 5-FOA resistant colonies (#1-6) were cultivated on medium lacking uridine (left, SC-URA) and medium containing 5 mM uridine (right, SC+Uri) for 7 days at 24°C.

**A**

| | |
|---|---|
| Wt | GCCAAGAAGCACTTCCTCGTCTTCGAGGACCGCAAGTTTGGCGACATTGGCAGTAGGTGCCTGCCCCACC |
| #1 | GCCAAGAAGCACTTCCTCGTCTTCGAGGACCGCAAGTTTG-----------------------CCCCACC |
| #2 | GCCAAGAAGCACTTCCTCGTCTTCGAGGACCGCAAGTTTGGCGA-ATTGGCAGTAGGTGCCTGCCCCACC |

**B**

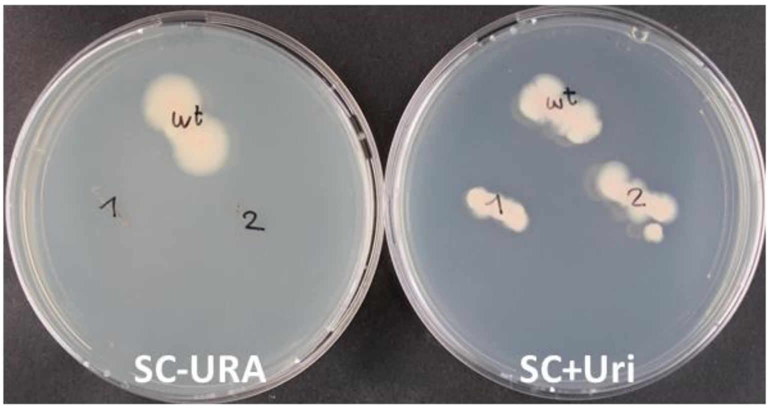


**Fig S6** CRISPR/Cas9 mediated manipulation of *ura3* in *A. pullulans* NBB 7.2.1 with RNP delivery.

**A** Partial sequences of *ura3* coding region of the *A. pullulans* NBB 7.2.1 wildtype (wt) and two 5-FOA resistant colonies (#1 and 2) obtained after the delivery of Cas9 RNPs. Target site of ura3_sgRNA1 is highlighted in green. PAM site is underlined. In all candidates deletions occurred at the sgRNA target site, resulting in a frame shift.

**B** The *A. pullulans* NBB 7.2.1 wildtype (wt) and the two 5-FOA resistant colonies (#1 and 2) were cultivated on medium lacking uridine (left, SC-URA) and medium containing 5 mM uridine (right, SC+Uri) for 7 days at 24°C.

GGCGACTCAATATGCATGCTCAAGACACACGCAGACATTATCAATGACTTTGGCCCTCGCACCATTCAAGGCCTG
AAAGAGATTGCCGCCAAAAAACACTTCCTCGTTTTCGAGGA**T**CGCAAGTTTGGCGACATTGGCAGTATGTTCATG
CCCCACTATCGTCTTGCTCGAACCACACACTGACAGCTGCTCGCAGGCACGGTGCAGAAGCAGTTCACCGCTGGG
CCTCTGCAAATCGTCCGCTGGGCAAACATTATCAACGCTCACATCTTTCCTGGCCCTGCCATCATTACTGCTCTC
TCGCAAGCTGCCCACGACGCCGTTACTTCGCTCAATACCGCAGTGACCACTTCCATCTCAGCCTCACCCGTTCCT
TCGTACATGGATGACAGCGATGAGGTCGACTCTCCTGCCTTGTCTCACCACGACGATTCGGATGACCTTGACAGG
ATGTCCAGCGACGAGGATACCAACCAGCTGTCTACATACAACGACCCTACCGGTCGCAAGCCCAGTGTCGTTTCC
GTCTCTACCACCATCAGCACAAAGACAGAAAGCATCTCACCTCAACCCACACCGAACCACCTTGGTGGTCCTTCC
GACTCCATCTCGGCTATCAGTGCAGGCTCGGCTAGTCAAGAATCCTCTGCTTTGGCCCGCCTCGGCGAACCTCCT
CTCCTTCGAAGCTTGCTCATTCTTGCAGAGATGAGCAGCGCGGGTAACTTGATGACTGGTGCATATACGGAACAA
TGTGTAGTTGAAGCACGCAAAAACCCGGAGTTTGTCATGGGCTTCAT

**Fig. S7** Sequence of part of the *ura3* gene in *A. pullulans* ATCC 42023. Primer ura3_2_fwd is indicated in blue, primer ura3_2_rev is indicated in green and ura3_sgRNA1 target site is indicated in yellow (mismatched base is indicated in bold).

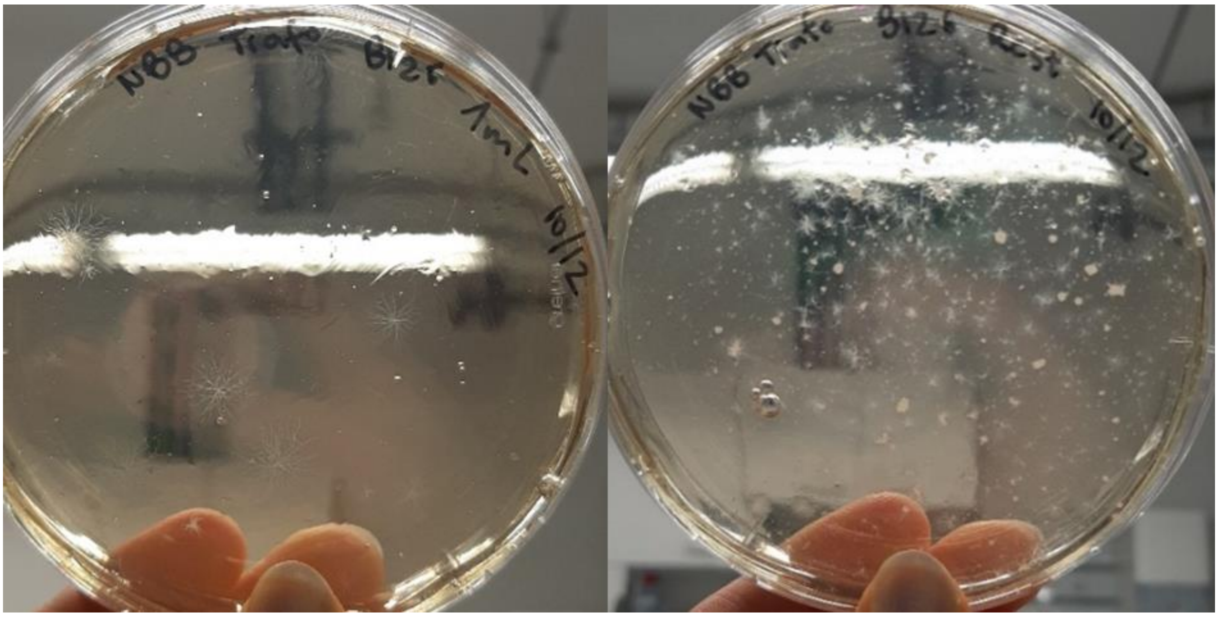

**Fig. S8** Selection plate (MEX + Uri + 5-FOA) resulting from the delivery of ura3_sgRNA1-Cas9 RNPs into *A. pullulans* NBB 7.2.1 with addition of β-mercaptoethanol to protoplasts and sgRNA denaturing and refolding before RNP assembly, after 12 days. About 50 CFUs were obtained after the delivery of 0.084 nmol Cas9 and sgRNA each.

```
        L  G  S  R  P  L  Y  A  E  K  W  A  N  F  K  A  E  L  A  V  M  V  V  K  T  K
Wt  514 CTTGGAAGCCGTCCTTTGTATGCTGAGAAGTGGGCCAACTTCAAGGCTGAGTTGGCAGTTATGGTTGTTAAGACCAAG 591

        L  G  S  R  P  L  Y  A  E  K  W  A  N  L                 V  M  V  V  K  T  K
#10 514 CTTGGAAGCCGTCCTTTGTATGCTGAGAAGTGGGCCAACTT---------------AGTTATGGTTGTTAAGACCAAG 576

        L  G  S  R  P  L  *
#13 514 CTTGGAAGCCGTCCTTTGTA-------------------------------------GTTATGGTTGTTAAGACCAAG 554

        L  G  S  R  P  L  Y  A  E  K  W  A  N  F  K  A         Q  L  W  L  L  R  P
#17 514 CTTGGAAGCCGTCCTTTGTATGCTGAGAAGTGGGCCAACTTCAAGGCT-------CAGTTATGGTTGTTAAGACCAAG 584

        L  G  S  R  P  L  Y  A  E  K  W  A  N  L                 V  M  V  V  K  T  K
#21 514 CTTGGAAGCCGTCCTTTGTATGCTGAGAAGTGGGCCAACTT---------------AGTTATGGTTGTTAAGACCAAG 576
```

**Fig S9** CRISPR/Cas9 mediated manipulation of *praics* in *A. pullulans* EXF-150 via co-delivery of RNPs targeting the *ura3* and the *praics* genes.

Partial sequences of *praics* coding region of the *A. pullulans* EXF-150 wildtype (wt) and the four adenine auxotrophic mutants (#10, 13, 17 and 21) obtained after the delivery of Cas9 RNPs. The corresponding amino acid sequence is given above the genomic sequence. Target site of praics_sgRNA is highlighted in yellow. PAM site is underlined. In all mutants deletions occurred at the sgRNA target site. In mutant #13 this lead to an early stop codon (*). In mutant #21 deletion of nucleotides resulted in a frame shift.

```
         M  P  Q  K  K  N  P  D  S  L  E  L  I  R  G  K  S  G  R  A  Y  G  Q  M  A
Wt  1025 ATGCCCCAAAAGAAGAACCCCGACTCGCTAGAACTCATCCGCGGAAAGTCTGGCAGAGCATACGGTCAGATGGCA 1099

         M  P  Q  K  K  N  P  D  S  L  E  L  K         S  L  A  E  H  T  V  R  W
#21 1025 ATGCCCCAAAAGAAGAACCCCGACTCGCTAGAACTCA-------AAAGTCTGGCAGAGCATACGGTCAGATGGCA 1092
```

**Fig. S10** CRISPR/Cas9 mediated manipulation of *asl* in *A. pullulans* EXF-150 via co-delivery of RNPs targeting the *ura3* and the *asl* genes.

Partial sequences of *asl* coding region of the *A. pullulans* EXF-150 wildtype (wt) and the arginine auxotrophic mutant (#21) obtained after the delivery of Cas9 RNPs. The corresponding amino acid sequence is given above the genomic sequence. Target site of asl_sgRNA is highlighted in yellow. PAM site is underlined. Deletions occurred at the sgRNA target site, resulting in a frame shift.

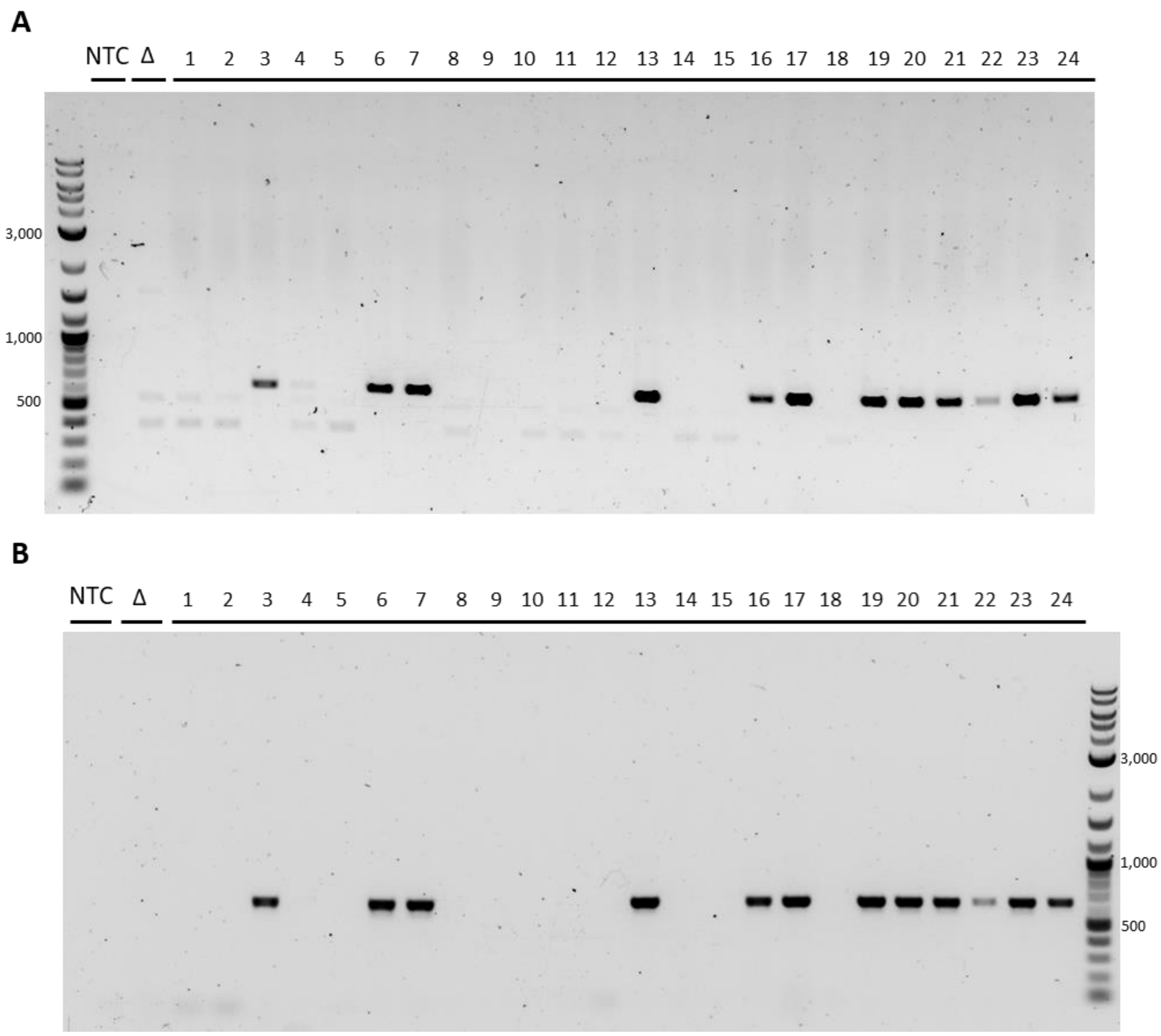


**Fig. S11** Agarose gel electrophoresis of the fragments obtained by PCR to test for homologous integration at the target locus in 24 candidates (#1-24) obtained after transformation of 3µg donor DNA with 20 bp-long homology flanks, without sgRNA-Cas9 RNPs. HR frequency = 12/24 = 50%.

**A** Agarose gel electrophoresis of the fragments obtained by PCR with primers dl4_20test_fwd (2) and pyr4_flank_rev (3) (Fig. 4) using chromosomal DNA of the parent strain (Δura3 #6, Δ) and 24 candidates as template. NTC, no template control.

PCR product of candidates # 3, 6, 7, 13, 16, 17, 19-24 is of expected length (618 bp).

**B** Agarose gel electrophoresis of the fragments obtained by PCR with primers dl4_20test_rev (5) and pyr4_flank_rev (6) (Fig. 4) using chromosomal DNA of the parent strain (Δura3 #6, Δ) and 24 candidates as template. NTC, no template control.

PCR product of candidates # 3, 6, 7, 13, 16, 17, 19-24 is of expected length (638 bp).

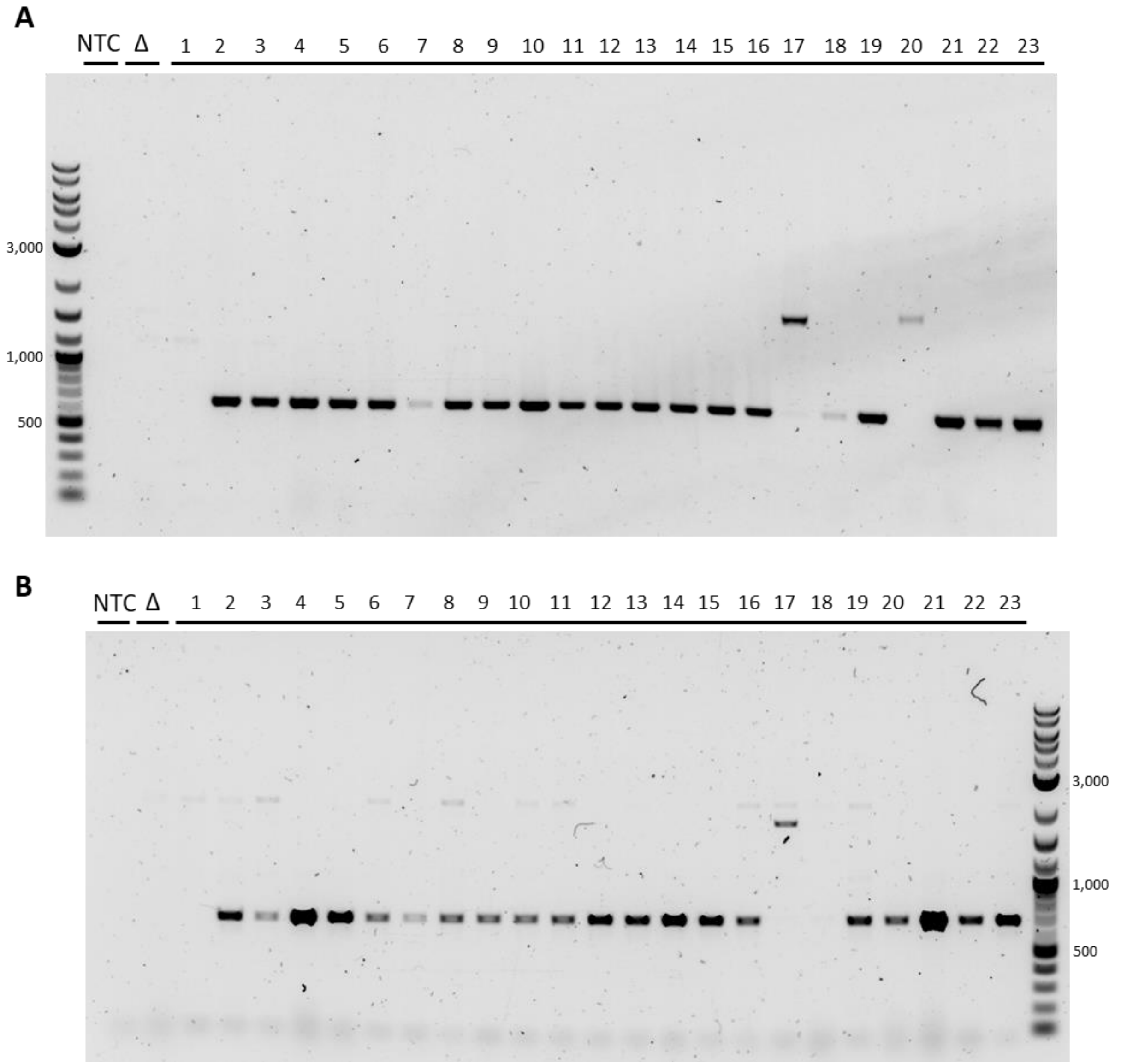


**Fig. S12** Agarose gel electrophoresis of the fragments obtained by PCR to test for homologous integration at the target locus in 23 candidates (#1-23) obtained after transformation of 3µg donor DNA with 500 bp-long homology flanks, without sgRNA-Cas9 RNPs. HR frequency = 19/23 =83%.

**A** Agarose gel electrophoresis of the fragments obtained by PCR with primers dl4_500test_fwd (1) and pyr4_flank_rev (3) (Fig. 4) using chromosomal DNA of the parent strain (Δura3 #6, Δ) and 23 candidates as template. NTC, no template control.

PCR product of candidates # 2-16, 18, 19, 21-23 is of expected length (643 bp).

**B** Agarose gel electrophoresis of the fragments obtained by PCR with primers dl4_500test_rev (4) and pyr4_flank_rev (6) (Fig. 4) using chromosomal DNA of the parent strain (Δura3 #6, Δ) and 23 candidates as template. NTC, no template control.

PCR product of candidates # 2-16, 19-23 is of expected length (667 bp).

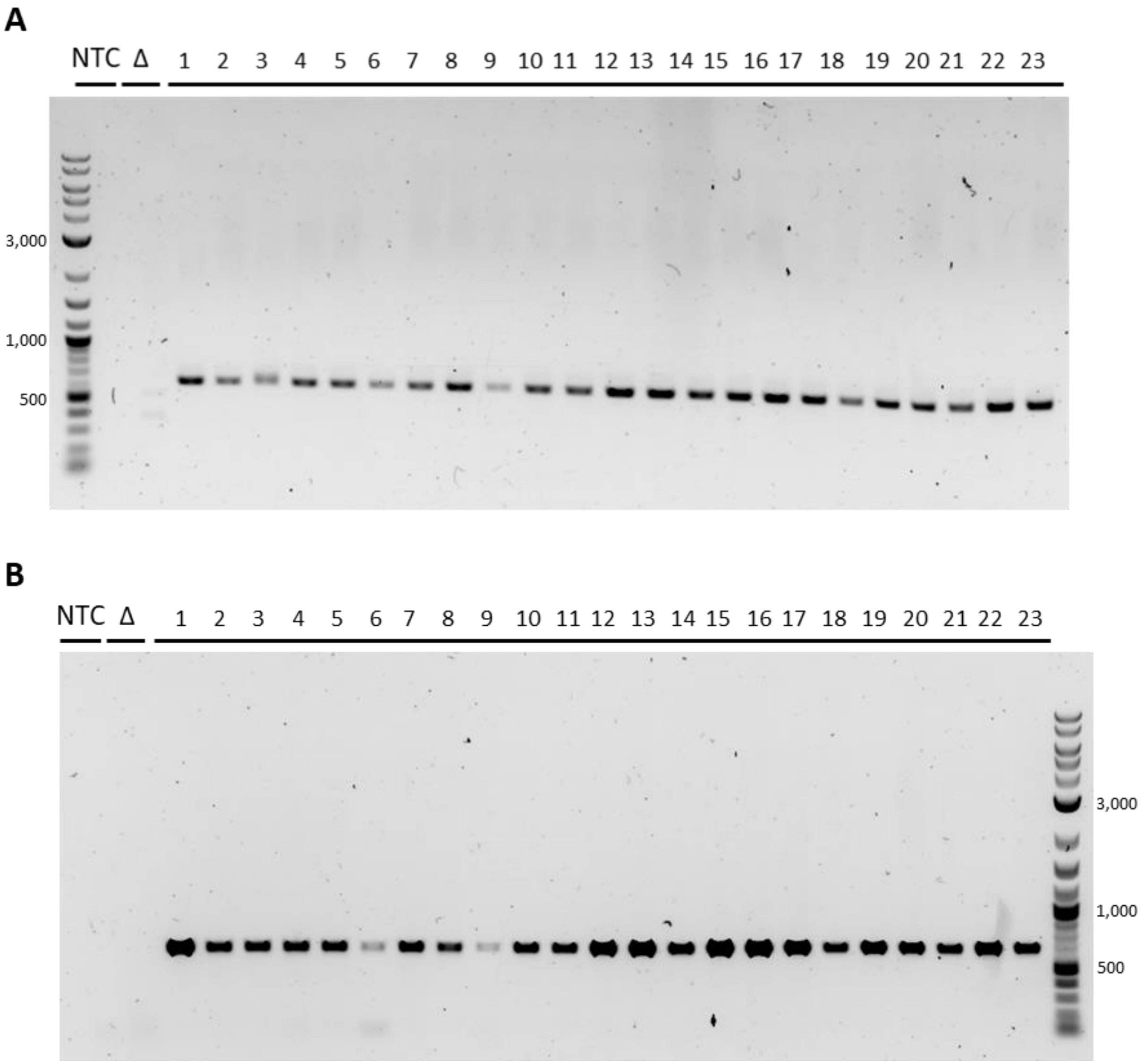


**Fig. S13** Agarose gel electrophoresis of the fragments obtained by PCR to test for homologous integration at the target locus in 23 candidates (#1-23) obtained after transformation of 3µg donor DNA with 20 bp-long homology flanks, with sgRNA-Cas9 RNPs. HR frequency = 22/23 = 96%.

**A** Agarose gel electrophoresis of the fragments obtained by PCR with primers dl4_20test_fwd (2) and pyr4_flank_rev (3) (Fig. 4) using chromosomal DNA of the parent strain (Δura3 #6, Δ) and 23 candidates as template. NTC, no template control.

PCR product of candidates # 1, 2, 4-23 is of expected length (618 bp). PCR product of #3 is slightly longer than expected.

**B** Agarose gel electrophoresis of the fragments obtained by PCR with primers dl4_20test_rev (5) and pyr4_flank_rev (6) (Fig. 4) using chromosomal DNA of the parent strain (Δura3 #6, Δ) and 23 candidates as template. NTC, no template control.

PCR product of all candidates is of expected length (638 bp).

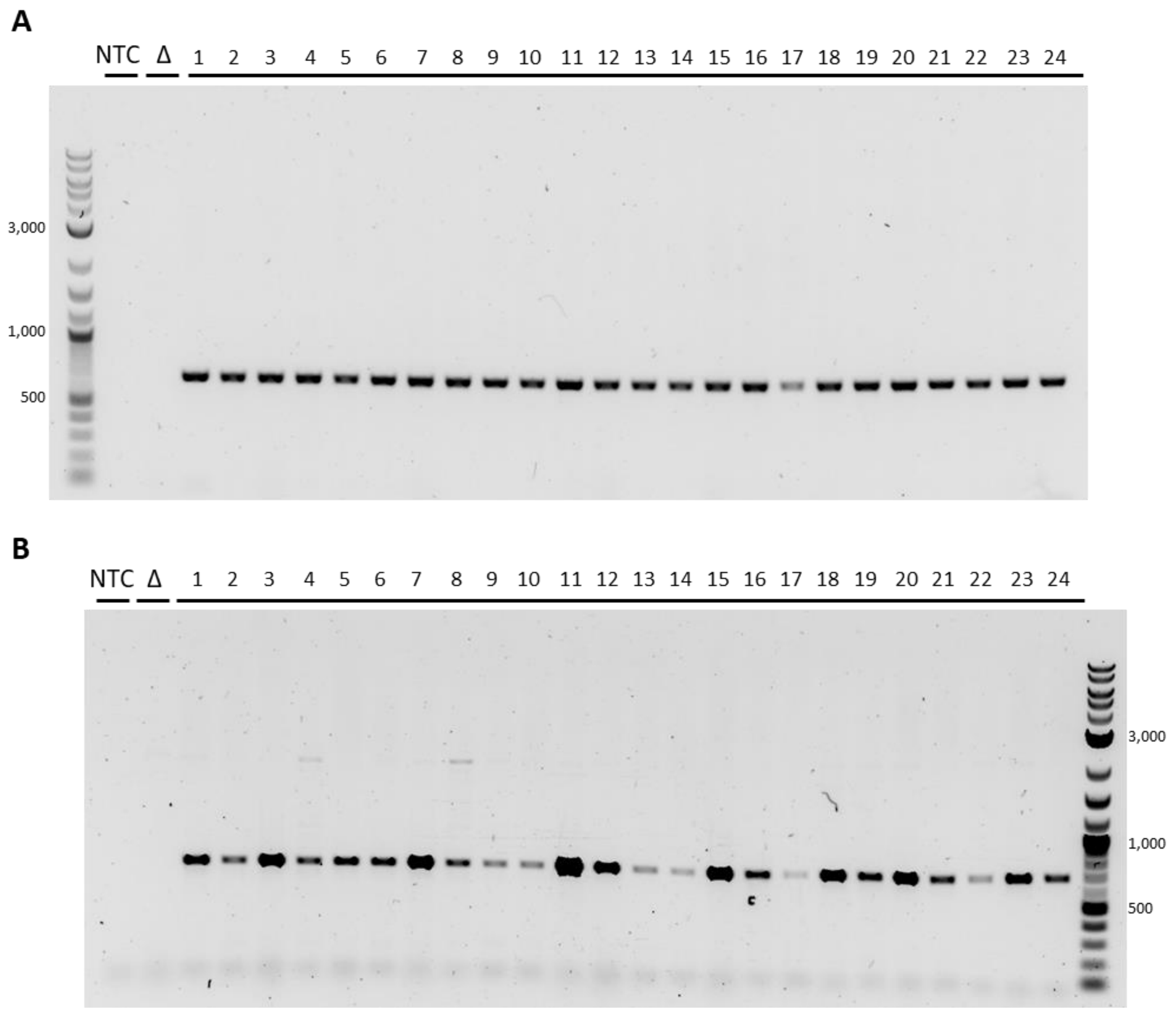


**Fig. S14** Agarose gel electrophoresis of the fragments obtained by PCR to test for homologous integration at the target locus in 24 candidates (#1-24) obtained after transformation of 3µg donor DNA with 500 bp-long homology flanks, with sgRNA-Cas9 RNPs. HR frequency = 24/24 = 100%.

**A** Agarose gel electrophoresis of the fragments obtained by PCR with primers dl4_500test_fwd (1) and pyr4_flank_rev (3) (Fig. 4) using chromosomal DNA of the parent strain (Δura3 #6, Δ) and 24 candidates as template. NTC, no template control.

PCR product of all candidates is of expected length (643 bp).

**B** Agarose gel electrophoresis of the fragments obtained by PCR with primers dl4_500test_rev (4) and pyr4_flank_rev (6) (Fig. 4) using chromosomal DNA of the parent strain (Δura3 #6, Δ) and 24 candidates as template. NTC, no template control.

PCR product of all candidates is of expected length (667 bp).

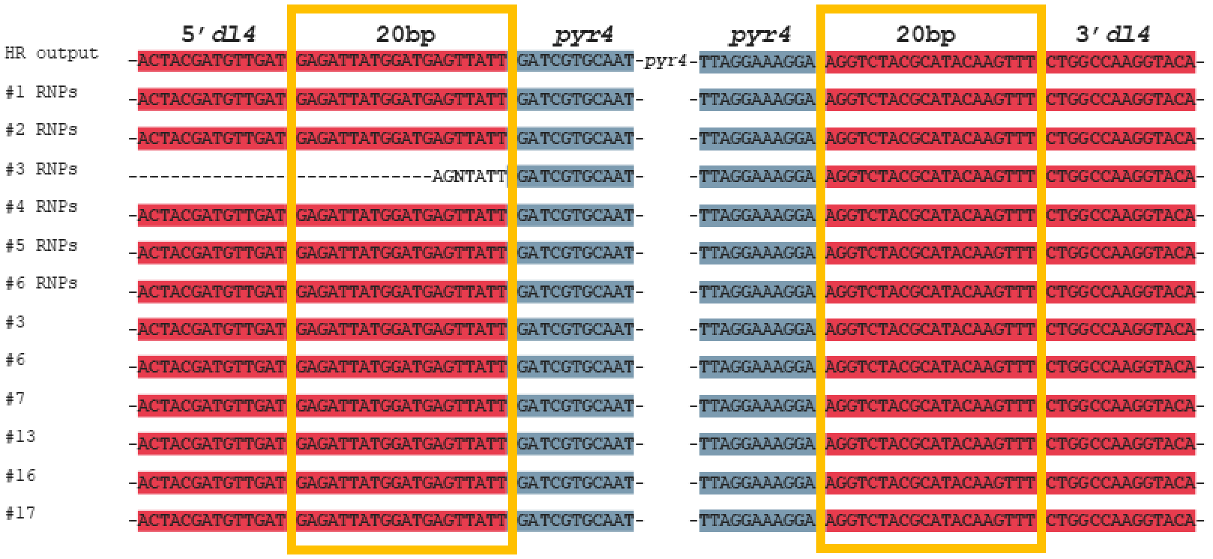


**Fig. S15** Sequence of the *dl4* insertion site in the theoretical HR output and 12 colonies obtained after transformation of donor DNA with 20 bp-long homology flanks (six with RNPs, #1, 2, 3, 4, 5 and 6; six without RNPs, #3, 6, 7, 13, 16 and 17). Homologous recombination could be confirmed in 11 transformants. One transformant (#3 RNPs) showed homologous recombination at the 3’flank but an inconclusive sequencing result at the 5’ site. Notably the corresponding PCR product is slightly longer than expected (Fig. S12). Red indicates *dl4* sequence; blue indicates *pyr4* sequence; yellow frame indicates 20bp-long homologous flank.

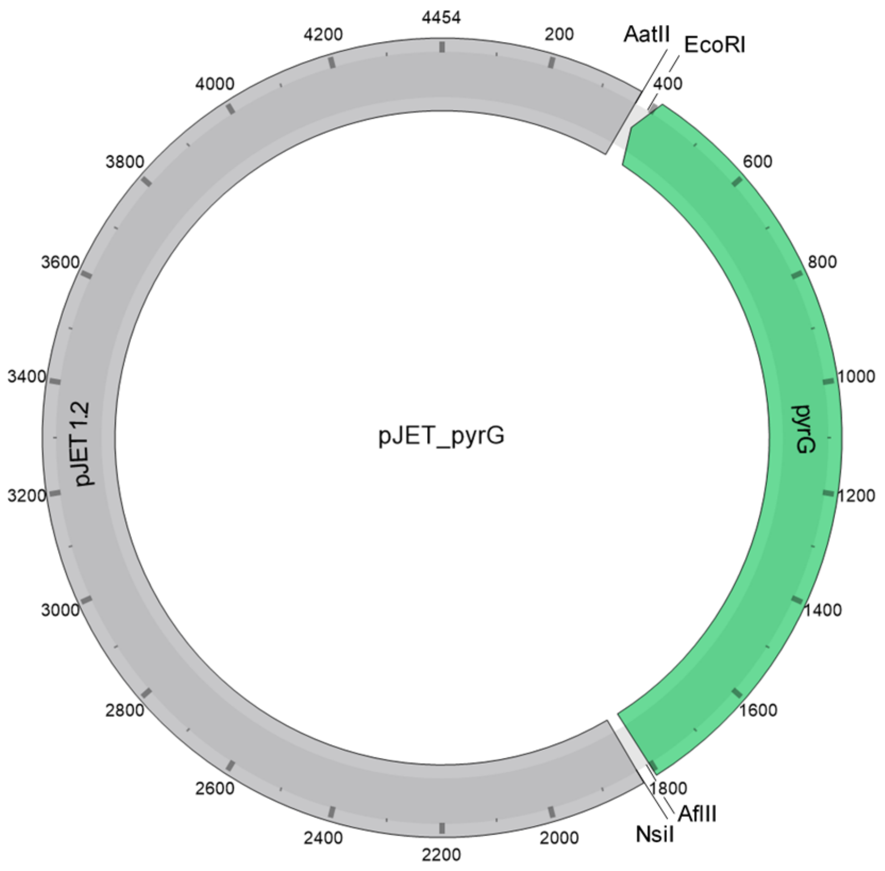


**Fig. S16 Vector map of pJET_pyrG**

GCCCCTGCAGCCGAATTATATTATTTTTGCCAAATAATTTTTAACAAAAGCTCTGAAGTCTTCTTCATTTAAATTCTTAGATG
ATACTTCATCTGGAAAATTGTCCCAATTAGTAGCATCACGCTGTGAGTAAGTTCTAAACCATTTTTTTATTGTTGTATTATCT
CTAATCTTACTACTCGATGAGTTTTCGGTATTATCTCTATTTTTAACTTGGAGCAGGTTCCATTCATTGTTTTTTTCATCATA
GTGAATAAAATCAACTGCTTTAACACTTGTGCCTGAACACCATATCCATCCGGCGTAATACGACTCACTATAGGGAGAGCGGC
CGCCAGATCTTCCGGATGGCTCGAGTTTTTCAGCAAGAT**GACGTC**TTTT**GAATTC**CTTGCTAGATGACTGGTAGGAATCTAAC
AAAGGGAAAACCTTGACGATTGGGGGCGGAAAACATGGCATTACAGGGGCCAATACGGACGAGCTCATCGTTTAATGACAGGC
ACAATGTATGCAGCCTTCCTCCACAACACTCGTACTGGCACTACACTTTTGCTCCAAGAGGAAATCATGACTTGCCGCATACT
CTGGCCATATAAGCTTCCCAGCCTTCTTTCTGGTACCGCTGTGCAGCTTCAACCGGGTCGGGAGCAGCGTAGATGCCTCGACC
GGCGATGATAAAGTCGGCACCGCGTCCAATAGCCGATGCAGGAGTCTGGTATTGCTGTCCAAGCTTATCTCCTTTGGAAGAGA
GGTTCACACCCGTCGTGAAGACCACGAAATCTTCATCCTCCGAGGCTGAAGACACATCCGACTGCACTTCCGTCAGGGCCCGC
GTCGACACGAAACCCATAACGAAGTTCTTGTATTTGCGAGCGTAGTCAACCGATGCCTTGGTATACTCGCCCGTAGCCAGCGA
TCCTTTGGGGGTCATCTCTGCCAGGACCAACAGTCCTCTCTCAGGACCATAGGGGAAGTCTTGCGCAGATGCGGTCTGGGCCA
GAGCCTCGACGATGCCCTCGCCAGGGAGAACGCTGCAGTTGATAATGTGGGCCCATTCGGAGATCCTCAGAGCACCGCCGTGG
TATTGCTTCTGGACGGTATTGCCGATGTCGATGAATTTGCGGTCCTCGAAGATCAAAAAGTTGTACTTTTGAGCCAGCACATT
CAGGCCATTGATAGTGTCGACGCTGAAATCGGTGAGGATGTCGATGTGTGTCTTGATGACGGCGATGTAGGGACCGAGACCTG
TATCATCAAAAGTTGATTAGTCCATTGCTAGACGGCATATGTATTGGATCCAACAGCTTCCGTACGGTCAGCGAGGTCCAGGA
GTTCTCGGGTTGTCGTCACATCAGCAGAGACGGTAACGTTTGTCTTCTTTGCTTCGGCAATCTCAAAAAGTCTCTTTGCCAGA
GGATTGGGGTGCTTGCTGGCTCGAGCACCGTAAGTCAATTGCGACTTGGACGACATCGTGGGAATGGAGGGTTATATGCGGGG
TATGACGGTGATTGATGAGTTGAAGTAGCCGAGCAATGAGGTATATTATCCAATTGAGGGTGACCCAACCAAATCAATTGCTT
GAGAATCAAATCTCAAATATTCTAAAATAGAAGCCGCCGGGGACTAAATTTAAAGTCGTACTCTATCTCGATCTTTTCCGCGT
GTAAAAATTGCGTGCCAATCTGGATGGAGACAGGCCACATCGGTGCTGTATTCCTCCGCCTCCGACTGTGGGGTCAACCACAA
TTAAACTACTCTGTAGTCAGGTACAGCTAGAATGGGGTAGACAGGCAGAACTGTAGAAGATAAAACATTGGTCAATCACTGGT
GGATCCTCTA**CTTAAG**AAA**ATGCAT**ATCTTTCTAGAAGATCTCCTACAATATTCTCAGCTGCCATGGAAAATCGATGTTCTTC
TTTTATTCTCTCAAGATTTTCAGGCTGTATATTAAAACTTATATTAAGAACTATGCTAACCACCTCATCAGGAACCGTTGTAG
GTGGCGTGGGTTTTCTTGGCAATCGACTCTCATGAAAACTACGAGCTAAATATTCAATATGTTCCTCTTGACCAACTTTATTC
TGCATTTTTTTTGAACGAGGTTTAGAGCAAGCTTCAGGAAACTGAGACAGGAATTTTATTAAAAATTTAAATTTTGAAGAAAG
TTCAGGGTTAATAGCATCCATTTTTTGCTTTGCAAGTTCCTCAGCATTCTTAACAAAAGACGTCTCTTTTGACATGTTTAAAG
TTTAAACCTCCTGTGTGAAATTATTATCCGCTCATAATTCCACACATTATACGAGCCGGAAGCATAAAGTGTAAAGCCTGGGG
TGCCTAATGAGTGAGCTAACTCACATTAATTGCGTTGCGCTCACTGCCAATTGCTTTCCAGTCGGGAAACCTGTCGTGCCAGC
TGCATTAATGAATCGGCCAACGCGCGGGGAGAGGCGGTTTGCGTATTGGGCGCTCTTCCGCTTCCTCGCTCACTGACTCGCTG
CGCTCGGTCGTTCGGCTGCGGCGAGCGGTATCAGCTCACTCAAAGGCGGTAATACGGTTATCCACAGAATCAGGGGATAACGC
AGGAAAGAACATGTGAGCAAAAGGCCAGCAAAAGGCCAGGAACCGTAAAAAGGCCGCGTTGCTGGCGTTTTTCCATAGGCTCC
GCCCCCCTGACGAGCATCACAAAAATCGACGCTCAAGTCAGAGGTGGCGAAACCCGACAGGACTATAAAGATACCAGGCGTTT
CCCCCTGGAAGCTCCCTCGTGCGCTCTCCTGTTCCGACCCTGCCGCTTACCGGATACCTGTCCGCCTTTCTCCCTTCGGGAAG
CGTGGCGCTTTCTCATAGCTCACGCTGTAGGTATCTCAGTTCGGTGTAGGTCGTTCGCTCCAAGCTGGGCTGTGTGCACGAAC
CCCCCGTTCAGCCCGACCGCTGCGCCTTATCCGGTAACTATCGTCTTGAGTCCAACCCGGTAAGACACGACTTATCGCCACTG
GCAGCAGCCACTGGTAACAGGATTAGCAGAGCGAGGTATGTAGGCGGTGCTACAGAGTTCTTGAAGTGGTGGCCTAACTACGG
CTACACTAGAAGGACAGTATTTGGTATCTGCGCTCTGCTGAAGCCAGTTACCTTCGGAAAAAGAGTTGGTAGCTCTTGATCCG
GCAAACAAACCACCGCTGGTAGCGGTGGTTTTTTTGTTTGCAAGCAGCAGATTACGCGCAGAAAAAAAGGATCTCAAGAAGAT
CCTTTGATCTTTTCTACGGGGTCTGACGCTCAGTGGAACGAAAACTCACGTTAAGGGATTTTGGTCATGAGATTATCAAAAAG
GATCTTCACCTAGATCCTTTTAAATTAAAAATGAAGTTTTAAATCAATCTAAAGTATATATGAGTAAACTTGGTCTGACAGTT
ACCAATGCTTAATCAGTGAGGCACCTATCTCAGCGATCTGTCTATTTCGTTCATCCATAGTTGCCTGACTCCCCGTCGTGTAG
ATAACTACGATACGGGAGGGCTTACCATCTGGCCCCAGTGCTGCAATGATACCGCGAGACCCACGCTCACCGGCTCCAGATTT
ATCAGCAATAAACCAGCCAGCCGGAAGGGCCGAGCGCAGAAGTGGTCCTGCAACTTTATCCGCCTCCATCCAGTCTATTAATT
GTTGCCGGGAAGCTAGAGTAAGTAGTTCGCCAGTTAATAGTTTGCGCAACGTTGTTGCCATTGCTACAGGCATCGTGGTGTCA
CGCTCGTCGTTTGGTATGGCTTCATTCAGCTCCGGTTCCCAACGATCAAGGCGAGTTACATGATCCCCCATGTTGTGCAAAAA
AGCGGTTAGCTCCTTCGGTCCTCCGATCGTTGTCAGAAGTAAGTTGGCCGCAGTGTTATCACTCATGGTTATGGCAGCACTGC
ATAATTCTCTTACTGTCATGCCATCCGTAAGATGCTTTTCTGTGACTGGTGAGTACTCAACCAAGTCATTCTGAGAATAGTGT
ATGCGGCGACCGAGTTGCTCTTGCCCGGCGTCAATACGGGATAATACCGCGCCACATAGCAGAACTTTAAAAGTGCTCATCAT
TGGAAAACGTTCTTCGGGGCGAAAACTCTCAAGGATCTTACCGCTGTTGAGATCCAGTTCGATGTAACCCACTCGTGCACCCA
ACTGATCTTCAGCATCTTTTACTTTCACCAGCGTTTCTGGGTGAGCAAAAACAGGAAGGCAAAATGCCGCAAAAAAGGGAATA
AGGGCGACACGGAAATGTTGAATACTCATACTCTTCCTTTTTCAATATTATTGAAGCATTTATCAGGGTTATTGTCTCATGAG
CGGATACATATTTGAATGTATTTAGAAAAATAAACAAATAGGGGTTCCGCGCACATTTCCCCGAAAGTGCCACCTGACGTCT
AAGAAACCATTATTATCATGACATTAACCTATAAAAATAGGCGTATCACGAGGCC

**Fig. S17 Sequence of pJET_pyrG**

The *pyrG* gene is indicated in green and the pJET backbone is indicated in grey. Bold letters indicate cut sites of restriction enzymes AatII, EcoRI, AflII and NsiI, respectively.

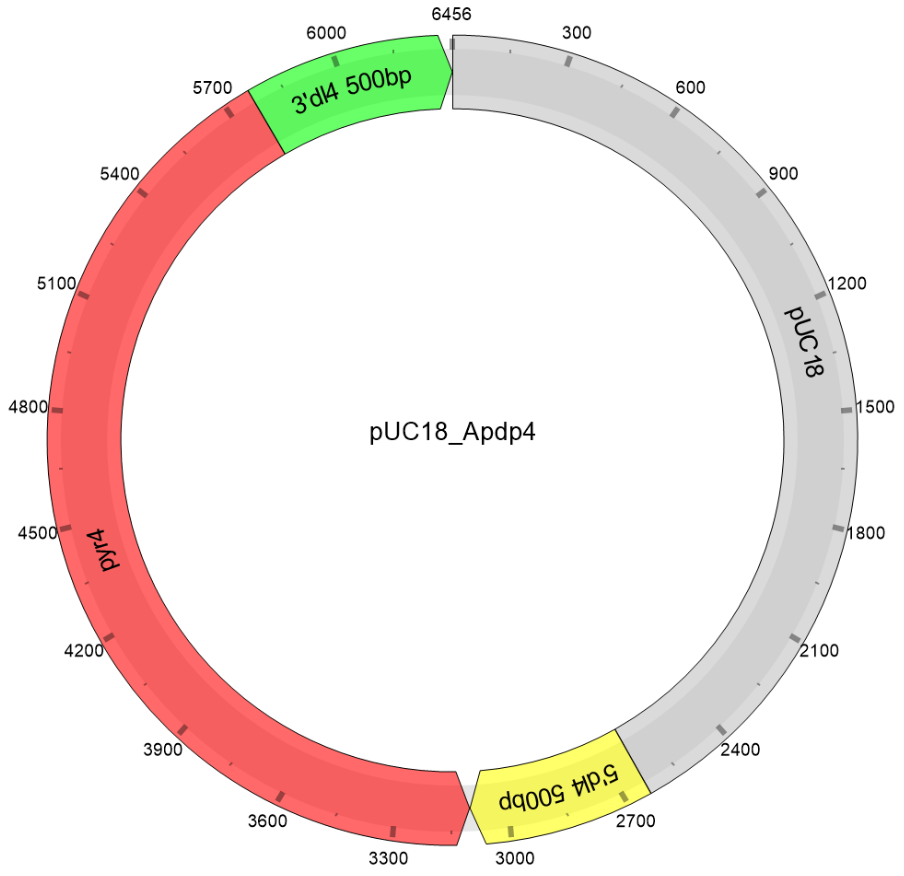


**Fig. S18 Vector map of pUC18_Apdp4**

ATGCAGGTCCTGAATCTGCTCTTTTCAACGGCATGAATTTCTACATCATCACTGGCGCTGCAAAGCCAATGAACAAGACCAAA
GCAGAATTGGAGCAACTGGTCAAGGCGAATGGAGGTAACATCGTCGCCACTCATAGTAATGCCGACACCATATGCATTGGAGA
AGGCAATCCCATTCGCATAGCTTCAATCAAGAAAGCTGGTACTCGAAACATCTTCAAGCCACACTGGTTGTTGGAATGCGTCA
AGCAGGCTGAAACAGATGTCGGAAGACCCAACGTCCTTCTCCCCTTCGAACCACGCCACGTCTTATTCAAGAAGGAGGAAGAC
GAGGACAGCTTCAACGGCAATACTGACGAATATGGCGACAGCTTCGCCCGAGATGTTGATGTTGAAGAATTGGAAAAGTTACT
TGCTGATATGCCCAAGTTCGAAGATGACGACTACGATGTTGATGAGATTATGGATGAGTTATTGATCGTGCAATTTCCTCTCC
TGGAGACATATCGCCTTGGTTGCCACCAGCGATCAATGGGTTTCTCACAGCACAAGTACTGCTTATCTGAATATTGTAGAGGG
GAGAACATGCCGCCGTGCTCTGAAGATGTTACGCTGCATACATTTAGAGGTACAGGCGCCATCACATGTCAATGTCACAAAGG
ATCGACACAAATAGCCATCAGGTAATACTCAATTGAACAATTCCGTATATATGGAAGCTGATATCGTCGACAACTGCATCCAA
ACCATCCTACCAAATGAAAAATGCCCGAGGGTGATATGTTCCCTCCACGCGCAAAAGCAAATGCCAAATCAAAAATACAATTC
CATGCTTCCAGATCCACCAGATATGTCAGGACCATATATGAACCTTCCAATGTGCAGTTCGCCTCCGCCATCGCGTGTTGTTG
CATGCAAAGATACACATCAATCGCAGCTGGGGTACAATCATCCATCATCCCAACTGGTACGTCATAACAAAAATCGACAAGAT
GGAAAAAGAGGTCGCCTAAATACAGCTGCATTCTATGATGCCGGGCTTTGGACAAGAGCTCTTTCTCAGCTCCGTTTGTCCTC
CCTCCCTTTTCCCCCTTCTTGCTAAATGCCTTTCTTTACTTCTTTCTTCCCTTCCCTCCCCTATCGCAGCAGCCTCTCGGTGT
AGGCTTTCCACGCTGCTGATCGGTACCGCTCTGCCTCCTCTACGGGGTCTGAGGCCTTGAGGATGCCCCGGCCCACAATGGCA
ATGTCGCTGCCGGCGATGCCAATCAGCTTGTGCGGCGTGTTGTACTGCTGGCCCTGGCCGTCTCCACCGACCGATCCGTTGGT
CTGCTGGTCCTCGTCTTCGGGGGGCAGCTGGCAGCCGGGCGTCATGTGGATAAAGGCATCGTCGGGCTCGGTGTTGAGCGTCT
CCTGCGAGATGAAGCCCATGACAAAGTCCTTGTGCTCCCGGGCGGCCTCGACGCAGGCCTGCGTGTACTCCTTGTTCATGAAG
TTGCCCTGGCTGGACATTTGGGCGAGGATCAGGAGGCCGCGGCTCAGCGGCGCCTCCTCGATGCCCGGGAAGAGCGACTCGTC
GCCCTCGGCGATGGCCTTTGTTAACCGGGGCGAGGAGACGGACTCGTACTGCTGGGTGACGGTGGTGATGGAGACGATGCTGC
CCTTGCGGCCGTCGCCGGACCGGTTCGAGTAGATGGGCTTGTCCAGGACGCCAATGGAGCCCATGCCGTTGACGGCGCCGGCG
GGCTCGGCGTCCCTGGAGTCGGCGTCGTCGTCAAACGAGTCCATGGTGGGCGTGCCGACGGTGACGGACGTCTTGACCTCGCA
GGGGTAGCGCTCGAGCCAGCGCTTGGCGCCCTGGGCCAGCGAGGCCACCGACGCCTTGCCGGGCACCATGTTGACGTTGACAA
TGTGCGCCCAGTCGATGATGCGCGCCGACCCGCCCGTGTACTGCAGCTCGACGGTGTGGCCAATGTCGCCAAACTTGCGGTCC
TCGAAGATGAGGAAGCCGTGCTTGCGCGCCAGCGACGCCAGCTGGGCTCCCGTGCCCGTCTCCGGGTGGAAGTCCCAGCCCGA
GACCATGTCGTAGTGCGTCTTGAGCACGACAATCGACGGGCCAATCTTGTCGGCCAGGTACAGCAGCTCGCGCGCTGTCGGCA
CGTCGGCGCTCAGGCACAGGTTGGACGCCTTGAGGTCCATGAGCTTGAACAGGTAAGCCGTCAGCGGGTGCGTCGCCGTCTCG
CTCCTGGCCGCGAAGGTGGCCTTGAGCGTCGGGTGTGGTGCCATGGCTGATGAGGCTGAGAGAGGCTGAGGCTGCGGCTGGTT
GGATAGTTTAACCCTTAGGGTGCCGTTGTGGCGGTTTAGAGGGGGGGAAAAAAAAGAGAGAGATGGCACAATTCTGCTGTGCG
AATGACGTTGGAAGCGCGACAGCCGTGCGGGAGGAAGAGGAGTAGGAACTGTCGGCGATTGGGAGAATTTCGTGCGATCCGAG
TCGTCTCGAGGCGAGGGAGTTGCTTTAATGTCGGGCTCGTCCCCTGGTCAAAATTCTAGGGAGCAGCGCTGGCAACGAGAGCA
GAGCAGCAGTAGTCGATGCTAGCGGAGAAGGCGATTCAATGCCGGCGGCGATACCCGATATTGCGACTTTGGGGGTGAGAAAA
AAAAAAAAAAAAAAACGGCGACCACCGAATGGGTTTCAGCGGCAGAGGTTTGGGTGCGGGAGGGTTGGTGGACCTGAGGAGCC
GAGAGGGTAGTAATGAGGGAGACGAGAGTGAGGATATGGAGGAGGAGGAGAAAGAAGAAGAAGAGGAAGAGGAGAAAAAGAGA
GAATAAAAGTAAAGAAAGGAATAAAAAAAAAGAAAAGGAAAGAAAATGAATTGCGCTGGACAAGGACTGAGGATGTTCGGCTC
AGGTCGCTCAGCAGCTGGAAGTGAAGCCCACCGGGTCCAGAAAGGGAGCTCGGTCAGGTCAAGTTTGAGGAGCCCTAGAAAGA
CGGACAACAGCTGGCAAACGTGACCAACGCCACAAAAAAATAAGAGTCGTAAATAGCGGTTGAGCCGTTCAGGTACAGTACAT
GTACTATGAAGCAGAAATAGGGTGAATGATACACACAAGTCTGCCAGATAATACCTACCCAGTCAGACCTTTTTCGTCAGCCA
AGATGGAGATGACAAGAAAAGGAAAAAGAGAAAGGTAGGGAAGTGGTTAGGAAAGGAAGGTCTACGCATACAAGTTTCTGGCC
AAGGTACACACATCGAGGCTGCGAAGCGTGTTGTTCTATTTGCCGGAGCACGTCTGGCGGACTCAATGGACGACGAGAAAATT
ACACATGTTGTGGCCGGATCAGAAGCGGAAGCAAGAGAATTGAGGATACAGACGGCCCGCCGCCGATATCCACCACGCGTTGT
CACAACAGGCTGGGTCATGAAGTCACTCAGAGAAGGCACGAGGCTTGATGAGGAGAGTGAGCTTTGAAAGCAAATAATGACCT
TGATGAACAATCGTCGGAGTTTGGCGGAGATAGCGACAGAAAACAAACGCCGCGGCATTTCATCATACCATATCTTCGCCATC
CACTATCCATATAAAATCCACAGAGTTTCAGTTCAAAAGCCCTCTTCCTTCAGCCAACCAGAAATCCGTCTCGATTCACAGCC
ATGTCTTCCAAAACTCTCATCCTCGTCACCGGCGCAAACCAAGGTCTCGGCTACTACGCCGCTCAACAACTCGCCGCTACTGG
CAATCATCACGTCCTCATTG

**Fig. S19 Sequence of linear disruption cassettes.**

The 5’ 500 bp-long homology flank is indicated in yellow and the 5’ 20 bp-long homology flank is indicated in blue. The *pyr4* gene is indicated in red. The 3’ 20 bp-long homology flank is indicated in orange and the 3’ 500 bp-long homology flank is indicated in green.

### Table S1: Oligonucleotides

| Name | Sequence (5’ – 3') |
|---|---|
| **CRISPR/Cas9 genome editing** | |
| S. pyogenes Cas9 scaffold oligo | AAAAGCACCGACTCGGTGCCACTTTTTCAAGTTGATAACGGACTAGCCTTATTTTAACTTGCTATTTCTAGCTCTAAAAC |
| ura3 specific DNA oligo 1 | TTCTAATACGACTCACTATAGACCGCAAGTTTGGCGACATGTTTTAGAGCTAGA |
| ura3_sgRNA1 | GACCGCAAGUUUGGCGACAUGUUUUAGAGCUAGAAAUAGCAAGUUAAAAUAAGGCUAGUCCGUUAUCAACUUGAAAAAGUGGCACCGAGUCGGUGCUUUU |
| ura3 specific DNA oligo 2 | TTCTAATACGACTCACTATAGGCACCGACATCATCATTGTGTTTTAGAGCTAGA |
| ura3_sgRNA2 | GGCACCGACAUCAUCAUUGUGUUUUAGAGCUAGAAAUAGCAAGUUAAAAUAAGGCUAGUCCGUUAUCAACUUGAAAAAGUGGCACCGAGUCGGUGCUUUU |
| praics specific DNA oligo | TTCTAATACGACTCACTATAGGCCAACTTCAAGGCTGAGTGTTTTAGAGCTAGA |
| praics_sgRNA | GGCCAACUUCAAGGCUGAGUGUUUUAGAGCUAGAAAUAGCAAGUUAAAAUAAGGCUAGUCCGUUAUCAACUUGAAAAAGUGGCACCGAGUCGGUGCUUUU |
| asl specific DNA oligo | TTCTAATACGACTCACTATAGACTCGCTAGAACTCATCCGGTTTTAGAGCTAGA |
| asl_sgRNA | GACUCGCUAGAACUCAUCCGGUUUUAGAGCUAGAAAUAGCAAGUUAAAAUAAGGCUAGUCCGUUAUCAACUUGAAAAAGUGGCACCGAGUCGGUGCUUUU |
| dl4 specific DNA oligo | TTCTAATACGACTCACTATAGGACCAGGATGTATGTTCAGGTTTTAGAGCTAGA |
| dl4_sgRNA | GGACCAGGAUGUAUGUUCAGGUUUUAGAGCUAGAAAUAGCAAGUUAAAAUAAGGCUAGUCCGUUAUCAACUUGAAAAAGUGGCACCGAGUCGGUGCUUUU |
| **Sequencing of *ura3* in EXF-150** | |
| ura3_fwd | ATGTCCCGTCACGCAACAATC |
| ura3_rev | TCACCTGCTACCTTTTGTTGC |
| **Sequencing of *ura3* in ATCC 42023 and NBB 7.2.1** | |
| ura3_2_fwd | GGCGACTCAATATGC |
| ura3_2_rev | ATGAAGCCCATGAC |
| **Sequencing of *praics*** | |
| praics_fwd | GTCACGCACAGAGGCATAC |
| praics_rev | GACGTCGTCGGTGATGTCG |
| **Sequencing of *asl*** | |
| asl_fwd | CTCAACCTCTAGAGATACTG |
| asl_rev | GGGTGATCATCTAACATAAC |
| **diagnostic PCR** | |
| pyrG_fwd | CTTGCTAGATGACTGGTAGG |
| pyrG_rev | ATGCATTTTTAGAGGATC |
| pyr4_fwd | ATATGAATTCGATCGTGCAATTTCCTCTCCTGG |
| pyr4_rev | ACTAGTGAGAGGCTGCTGCGATAGG |

| | |
|---|---|
| **Cloning** | |
| pyrG_fwd-AflII-NsiI | ATGCATTTTCTTAAGTAGAGGATCCACCAGTGATTG |
| pyrG_rev-EcoRI-AatII | GACGTCTTTTGAATTCCTTGCTAGATGACTGGTAGG |
| pyr4dl4_20bp_rev | AAACTTGTATGCGTAGACCTTCCTTTCCTAACCACTTCCCTACC |
| pyr4dl4_20bp_fwd | GAGATTATGGATGAGTTATTGATCGTGCAATTTCCTCTCCTGG |
| dl4_5Overlap_rev | AATAACTCATCCATAATCTCA |
| dl4_5Overlap500_fwd | GAGCTCGAATTCGTAATCATGATGCAGGTCCTGAATC |
| dl4_3Overlap_fwd | AGGTCTACGCATACAAGTTT |
| dl4_3Overlap500_rev | ACACAGGAAACAGCTATGACCAATGAGGACGTGATGATTG |
| pUC18_rev | CATGATTACGAATTCGAGCTC |
| pUC18_fwd | GTCATAGCTGTTTCCTGTGTGA |
| **preparative PCR** | |
| dl4_20bpover_rev | AAACTTGTATGCGTAGACCT |
| dl4_20bpover_fwd | GAGATTATGGATGAGTTATTGATC |
| dl4_500bpover_rev | CAATGAGGACGTGATGATTG |
| dl4_500bpover_fwd | ATGCAGGTCCTGAATC |
| **diagnostic PCR + sequencing** | |
| dl4_20test_fwd | ATGCAGGTCCTGAATC |
| dl4_500test_fwd | GCAGACGAACAGGTCAAAACG |
| pyr4_flank_rev | GCAGCGTAACATCTTCAGAGC |
| dl4_20test_rev | CAATGAGGACGTGATGATTG |
| dl4_500test_rev | CTTCTTAGCCTTGTTGACGTC |
| pyr4_flank_fwd | GATAATACCTACCCAGTCAGACC |
